# Generative AI designs functional thiolation domains for reprogramming non-ribosomal peptide synthetases

**DOI:** 10.64898/2026.03.03.709401

**Authors:** Emre F. Bülbül, Seounggun Bang, Kevin George, Gabriele Bianchi, Prateek Raj, Seonyong Chung, Vincent Pauline, Ramon Hochstrasser, Hannah A. Minas, Walid A. M. Elgaher, Andreas M. Kany, Anna K. H. Hirsch, Steven Schmitt, Dirk W. Heinz, Olga V. Kalinina, Dietrich Klakow, Kenan A. J. Bozhüyük

## Abstract

Large language models and generative protein design promise to accelerate biotechnology, but it remains unclear whether they can engineer dynamic megasynth(et)ases whose activity depends on transient, context-specific domain interfaces. Non-ribosomal peptide synthetases (NRPSs) are an especially demanding target, yet a high-value one because they produce many clinically important natural products and offer a route to analogs that are often difficult or impractical to access by chemical synthesis. Here we integrate pretrained generative models (ESM3, ProteinMPNN and EvoDiff) with design–build–test–learn cycles and data-guided prioritization to generate 76 *de novo* thiolation (T) domains. We built and tested 578 recombinant NRPS variants *in vivo* spanning minimal, full-length and hybrid assembly lines. AI-designed T-domains supported product formation across architectures, enabled catalytically active hybrids at recombined junctions and increased yields by up to ∼3-fold relative to NRPSs carrying the native T-domain. A representative design showed improved soluble expression, refolding, and a 12 °C higher melting temperature, while molecular dynamics simulations indicated preserved global stability but reshaped, state-dependent interdomain contact networks. Together, these results establish generative design as an effective route to context-conditioned optimization and reprogramming of biosynthetic assembly lines.

## Main

Protein engineering drives innovation across the life sciences but remains constrained by a fundamental search problem. Functional solutions are rare, sequence space is vast, and experiments are costly^1-3^. These limitations are particularly acute for multifunctional multidomain assembly line enzymes such as non-ribosomal peptide synthetases (NRPSs), where activity depends on coordinated interdomain dynamics^4,5^. Established approaches such as rational design and directed evolution are constrained by incomplete understanding of sequence-structure-function relationships and by the high experimental cost of exploring sequence space^6-8^.

These constraints have accelerated interest in artificial intelligence (AI) methods that learn from natural sequence diversity and can generate and rank candidate proteins before committing to costly experiments^9,10^. This has given rise to a rapidly expanding toolkit of generative approaches spanning several model classes. (I) Sequence-based foundation models such as ESM3^11^; (II) Diffusion models such as EvoDiff^12^; and (III) Structure-conditioned design methods such as ProteinMPNN^13^. Recent studies have demonstrated that protein language models can generate functional enzymes with native-like catalytic activity^14-16^. Whether such models can be extended to highly dynamic, multidomain systems remains unclear, particularly when function depends on coordinated conformational changes and state-dependent interdomain communication.

NRPSs represent an especially demanding setting for protein design^17^. They biosynthesize a vast diversity of bioactive natural products (NPs), including many clinically important antibiotics, immunosuppressants, and anticancer agents, through an assembly line mechanism in which modules sequentially extend and process a growing intermediate^18-22^. A canonical NRPS elongation module minimally comprises an adenylation (A) domain for substrate selection and activation, a thiolation (T) domain, also known as peptidyl carrier protein (PCP), which carries intermediates as thioesters on a 4’-phosphopantetheine (PPant) arm, and a condensation (C) domain that catalyzes peptide bond formation. In some modules, the condensation domain is bifunctional, forming a dual condensation/epimerization (C/E) domain that additionally catalyzes stereochemical inversion of the upstream residue^4^. Many NRPSs terminate in a thioesterase (TE) domain that releases the peptide either by hydrolysis (linear products) or by cyclization (macrocyclic products).

This modular architecture of NRPSs has inspired engineering efforts, ranging from reprogramming catalytic domains, to recombination of larger modular units aimed at generating NP analogs and entirely new-to-nature peptides^23-30^. In practice, however, functional outcomes remain difficult to predict, and many engineered systems lose activity^17,31^. A central reason is that NRPSs function as dynamic molecular machines rather than rigid assemblies, with T-domains cycling between multiple interaction partners during catalysis and forming transient interfaces that depend on the catalytic state^5,32-35^. Consequently, even subtle perturbations at domain junctions can disrupt interdomain communication and coordination, impairing productive turnover despite seemingly compatible module architectures.

Evolutionary patterns underscore these constraints. Comparative analyses suggest that recombination has contributed to NRPS diversification, but natural events tend to preserve partner-specific interdomain contacts rather than enabling unrestricted domain exchange^36^. This insight motivated evolution-guided engineering frameworks^37,38^ such as the eXchange Unit Thiolation (XUT) concept^39^, which defines fusion sites around and within T-domains that better maintain structural compatibility. This approach has enabled functional chimeras in several settings, including systems producing the antibiotic odilorhabdin and *de novo* designed proteasome inhibitors^39-41^. Yet, even with such guidance, engineered junctions frequently remain non-native. During catalysis, T-domains cycle through transient contacts with both upstream and downstream partners, and chimeric constructs inherently introduce non-cognate interfaces^42,43^. Consequently, recombination often yields functional compromises, typically reducing activity and in many cases abolishing production completely. These outcomes highlight that productive interdomain communication cannot be inferred from sequence motifs alone, but depends on context-specific interface features and dynamics.

These constraints raise the question of whether generative AI can design *de novo* T-domains that remain compatible with their A-T-C interface environment beyond evolutionary sequence space. Here, we address this challenge by coupling pretrained generative models with iterative design-build-test-learn (DBTL) cycles to create and experimentally validate new-to-nature T-domain sequences.

## Results

We established an AI-enabled computational and experimental workflow for *de novo* NRPS T-domain design and *in vivo* functional validation. Across successive DBTL rounds, we benchmarked AI-generated sequences against mutational controls and identified T-domain variants that supported catalytic activity across increasingly complex NRPS architectures, including minimal, full-length, repositioned, and chimeric assembly lines.

### A type S NRPS platform for high-throughput T-domain design and validation

To enable rapid, quantitative evaluation of AI-designed T-domains in an NRPS context, we used a bipartite type S variant of the GameXPeptide synthetase (GxpS)^44-48^, which produces GameXPeptides A-E, a family of cyclic nonribosomal peptides from *Photorhabdus luminescens* TTO1, a bacterium that lives in symbiosis with entomopathogenic nematodes^49,50^. The type S platform provides a two-tier *in vivo* activity readout (Fig. 1a): T-domain variants are first prescreened in the subunit 2 (SU2)-only configuration (A3–TE) by quantifying the two dominant linear tripeptides flL (**1**) and FlL (**2**) (Fig. 1a). As reported previously, the promiscuous A3-domain incorporates either phenylalanine or leucine, yielding a mixture of linear tripeptides (**1**–**4**), including D–D–L and L–D–L configured products^47^. Functional designs are then validated in the reconstituted full assembly line by coexpressing SU2 with the upstream SU1 (A1–C3), which yields macrocyclic pentapeptides (**5**–**8**); the main product cyclo(vLflL) (**6**) serves as the quantitative readout under these conditions (Fig. 1a). The type S architecture splits the native assembly line between the C3 and A3 domains to generate two independently expressed SUs that assemble *in vivo* via a synthetic zipper (SZ) interaction, a designed high-affinity coiled-coil interaction inspired by leucine zippers^47^ (Fig. 1b).

**Fig 1:**
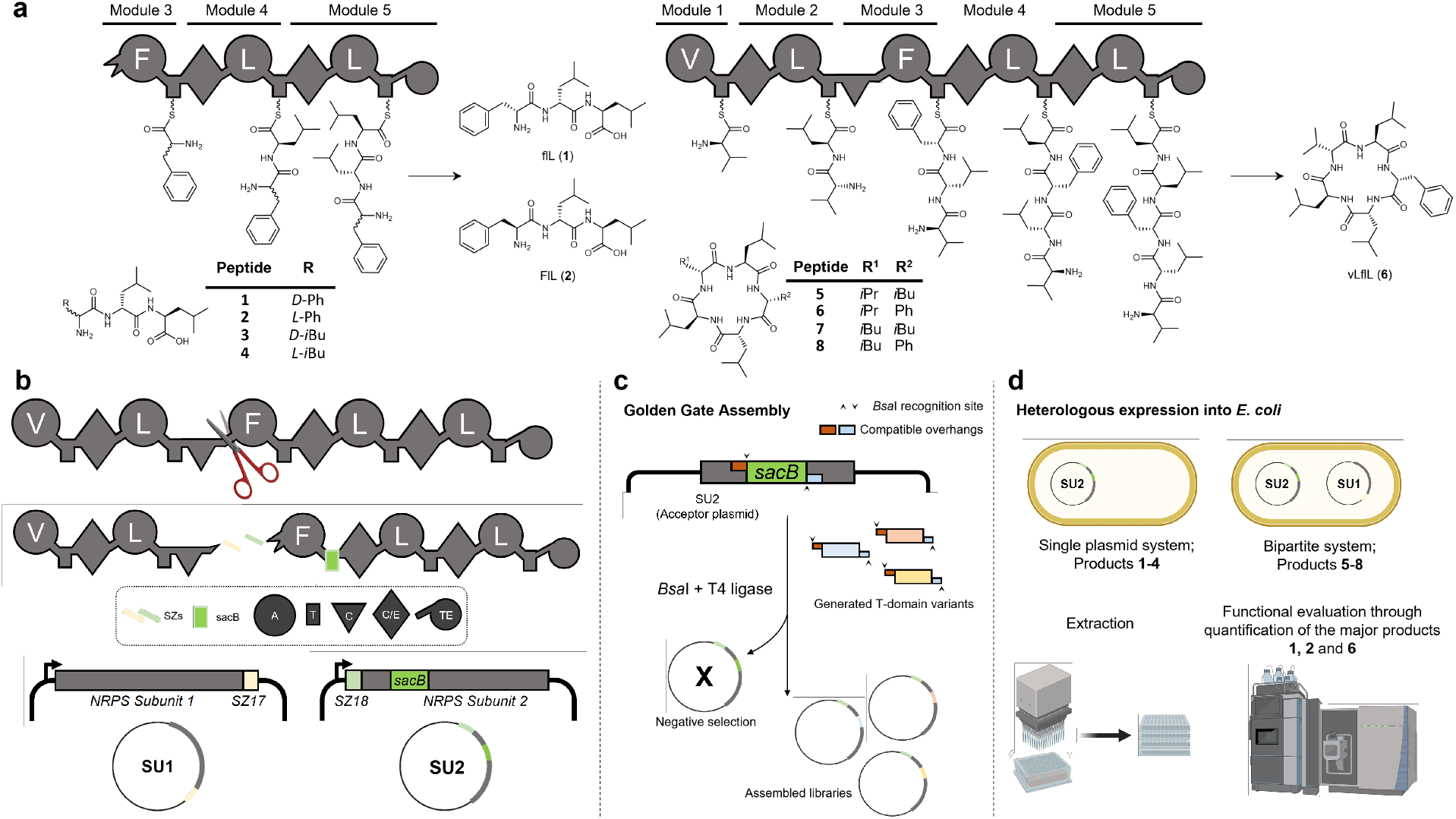
Type S GxpS architecture enables rapid exchange of T-domains via synthetic zippers^46^ and Golden Gate^51^ assembly. **a**, Schematic representation of truncated (left) and full-length wild-type GxpS (right) assembly line architectures. Chemical structures of the peptide products are shown, including the linear tripeptides **1** and **2** produced by the truncated construct and the macrocyclic pentapeptide product **6** produced by the full-length synthetase, as well as minor products. **b**, Schematic representation of native GxpS (top) and its bipartite type S counterpart (bottom). The assembly line is split between C3- and A3-domains, yielding two independently expressed NRPS protein subunits (SUs) that assemble *in vivo* via the synthetic zipper (SZ) pair SZ17 and SZ18. Each domain is represented by a distinct symbol. A, circle; T, rectangle; C, triangle; C/E, rhombus; and TE, tailed circle. Substrate specificities of the A-domains are indicated by capital letters following the standard amino acid code. **c**, High-throughput T-domain swapping in SU2 by Golden Gate replacement of BsaI-flanked *sacB* counterselection cassette with candidate T-domain sequences. **d**, Functional reconstitution of the split GxpS system in *E. coli*, illustrating the SU2-only prescreen and the full assembly line validation readouts.

To enable high-throughput swapping of T3 variants, we built a Golden Gate cloning^51^ workflow that replaces a BsaI-flanked sacB counterselection cassette in SU2 with candidate T-domain sequences (Fig. 1c). We then ran an integrated DBTL campaign that coupled conditional generative design to rapid *in vivo* validation (Fig. 1d). Candidate T-domains were proposed using three complementary generative strategies (ESM3^11^, EvoDiff^12^, ProteinMPNN^13^), under an explicit local-context constraint: we specified the intended A–T–C neighborhood by providing the flanking A and C domains (and, where applicable, structural context) while masking the T-domain segment during generation, thereby forcing designs to be compatible with the target junction environment (Supplementary Note 1; Supplementary Data 1). ProteinMPNN designs were conditioned on backbone coordinates, ESM3 designs leveraged structure tokens, and EvoDiff provided a sequence-only baseline. As the campaign progressed, we combined generative proposals with progressive candidate prioritization, using lightweight *in silico* filters and, in later rounds, data-driven surrogate models to focus testing on sequences most likely to be functional (details in Supplementary Note 1). Selected variants were cloned into the type S GxpS scaffold and quantified in *E. coli*, and the resulting sequence– function relationships informed two subsequent design rounds. Product identities for all GxpS-derived peptides were confirmed by tandem mass spectrometry (MS/MS) (Supplementary Fig. S1).

### AI-designed T-domains are frequently functional in a minimal NRPS assay system

We first benchmarked AI-designed T-domains in the SU2-only prescreen by quantifying production of **1** and **2** (Fig. 2). Across three iterative design rounds, we tested 66 AI-designed T-domains at the T3 position of the type S GxpS scaffold (Supplementary Data 1) and normalized activities to a wild-type (WT) T3 control included in each experiment (WT = 100 %). Quantification and product assignments were validated by targeted LC–MS/MS in selected reaction monitoring (SRM) using external standards (Supplementary Fig. S2; Supplementary Table S1).

**Fig 2:**
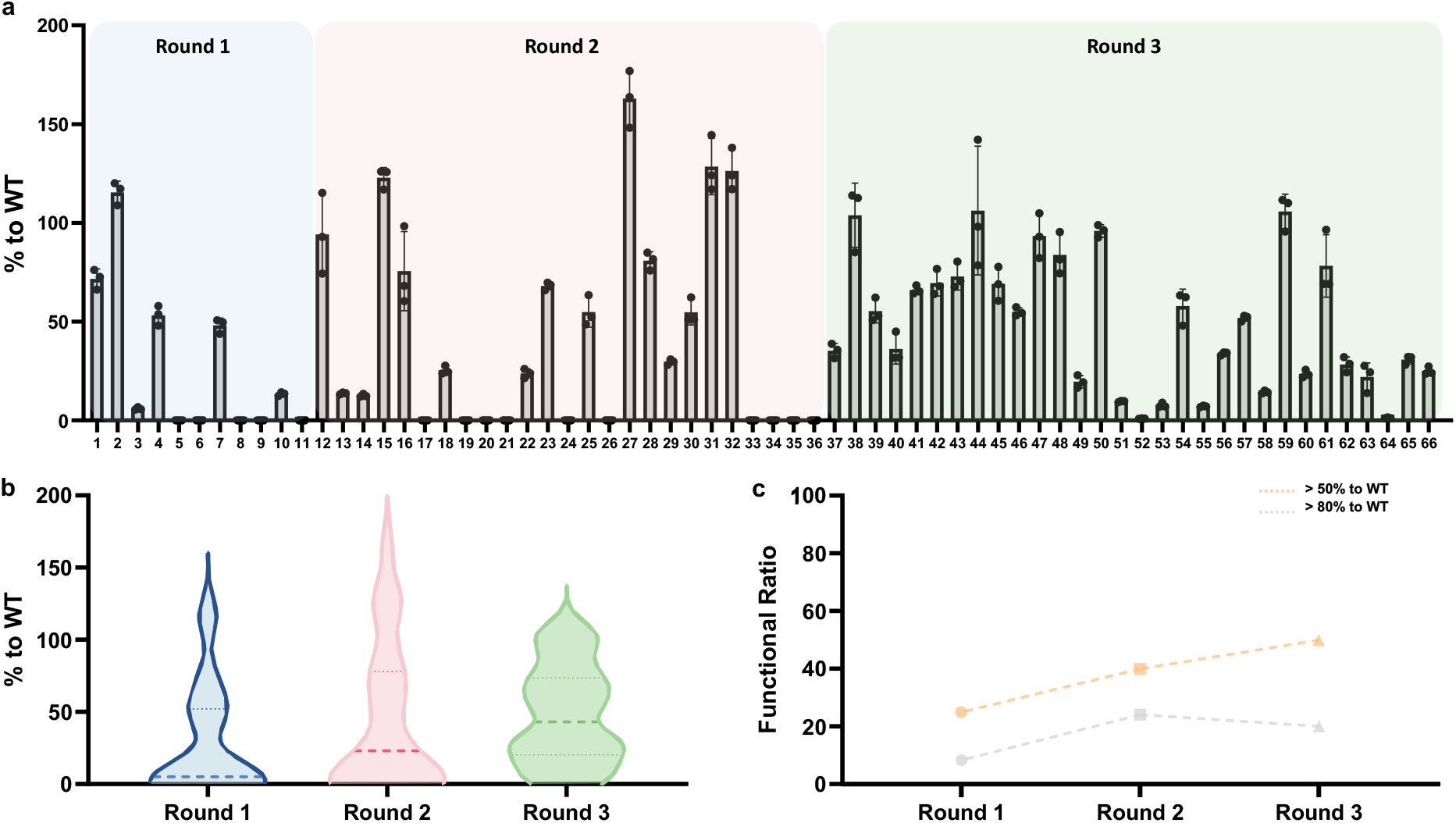
Round-by-round performance of AI-generated T-domain variants in the minimal system producing 1 and 2. **a**, Production of **1** and **2** for variants tested in round 1-3, expressed in *E. coli* and screened in succinate XPP medium at the 1 mL scale, shown relative to the WT GxpS_T3 control included in each experiment (WT = 100%). Bars show mean ± standard deviation (SD) (n = 3 biological replicates); points indicate individual replicates. Complete raw and processed datasets underlying all experiments are provided in Source Data and Supplementary data 1, respectively. Quantification was performed by LC-MS using external calibration with synthesized or purified standards (see Supplementary Fig. S2). **b**, Violin plots summarizing the distribution of activities for each round. **c**, Fraction of variants exceeding the indicated activity thresholds. Sample sizes: round 1 (n = 11), round 2 (n = 25), round 3 (n = 30).

In round 1 (Fig. 2a, see Supplementary Note 1 for details), we selected 11 variants (AI-1 to AI-11) from an initial generative pool produced with ESM3 and ProteinMPNN and assessed out-of-the-box performance without *in silico* filtering beyond sequence clustering and representative sampling. All four ESM3-derived variants (AI-1 to AI-4) supported detectable production of **1** and **2** when installed at T3, with AI-2 exceeding WT (∼115 %), whereas only two (AI-7 and AI-10) of seven ProteinMPNN-derived variants (AI-5 to AI-11) supported detectable production.

In round 2 (AI-12 to AI-26; Fig. 2a), we expanded the workflow to include EvoDiff and refined candidate selection using simple sequence-based filters informed by round 1 performance and *in silico* evaluation. To ensure post-translational activation of the T-domain by phosphopantetheinylation, we retained sequences containing the conserved FFxxGGxS motif that includes the active-site serine for 4′-phosphopantetheine attachment, imposed a minimum identity threshold of 50 % to WT GxpS_T3 and filtered for low perplexity using the ESM2 650M^52^ language model (4.8 or lower).

In round 3 (AI-27 to AI-66; Fig. 2a), candidate prioritization was further guided by surrogate models trained on the sequence–activity measurements obtained in rounds 1 and 2 together with an expanded set of error-prone polymerase chain reaction (epPCR)-derived variants described below (Supplementary Note 1). These models were used to rank newly generated sequences and select variants for experimental testing.

These refinements increased the frequency of functional designs across rounds (Fig. 2b, c). While the fraction of near-WT performers (≥80 %) peaked in round 2 and decreased in round 3, the proportion of variants exceeding a moderate activity threshold (≥50 %) increased steadily from round 1 to round 3, yielding a broader set of consistently functional T-domains. Sequence identity to WT GxpS_T3 had limited predictive value for NRPS output (Supplementary Fig. S3): variants spanning ∼50–75 % identity exhibited outcomes ranging from highly active to inactive. For example, incorporation of AI-27 at the T3 position increased production to ∼163 % relative to WT at ∼70.4 % identity to WT GxpS_T3, whereas AI-5 yielded no detectable product formation despite a comparable level of sequence similarity (∼64.3 % identity). Together, these results show that sequence identity to WT is insufficient to predict function in the SU2 prescreen, motivating subsequent tests of how junction context and partner identity shape compatibility in full-length and hybrid NRPS architectures.

### Benchmarking by error-prone mutagenesis reveals scaffold-dependent gains

After round 2, we used epPCR to benchmark classical local sequence exploration of WT GxpS_T3 and, in parallel, to diversify three AI-designed scaffolds (AI-2, AI-12 and AI-15) and expand the sequence–activity dataset for surrogate modeling. We screened the resulting libraries in the SU2-only assay, using production of **1** and **2** as the functional readout and normalizing activities to WT (WT = 100 %) (Supplementary Fig. S4a). Complete sequence lists for epPCR variants (IDs 77–258) and their corresponding production values are provided in Supplementary Data 1.

Across libraries, epPCR produced broad outcome distributions and revealed strong scaffold dependence (Supplementary Fig. S4a–c). In the WT GxpS_T3 library (n = 50), 12 variants (24 %) exceeded the WT control, whereas 8 variants (16 %) showed no detectable production. In the AI-12-derived library (parent output 94 % of WT; n = 42), only 3 variants (∼7 %) exceeded WT and 16 variants (∼38 %) were non-functional, with a maximum of 108 % (variant 192). In the AI-15-derived library (parent output 123 % of WT; n = 37), 7 variants exceeded WT and 8 were inactive, reaching up to 210 % (variant 147). By contrast, diversification of AI-2 (parent output 116 % of WT; n = 53) yielded a substantially higher fraction of improved variants: 32 variants (∼60 %) exceeded WT, 12 variants (∼23 %) were inactive. Notably, this library also produced the top-performing variant in the study (variant 123), reaching 285 % of WT.

Together, these data show that epPCR outcomes depend strongly on the starting scaffold. While the WT T3-domain sequence can be improved, AI-2 provided greater headroom for local refinement, yielding both a higher fraction of improved variants and larger gains in SU2 production of **1** and **2** (Supplementary Fig. S4c).

### AI-designed T-domains remain functional in the full-length type S GxpS system

After establishing that AI-designed T-domains support peptide biosynthesis in the SU2-only assay, we evaluated their performance in the reconstituted full-length type S GxpS assembly line, which comprises all five modules (Supplementary Fig. S1e). Peptide production was quantified using the main product **6**. Product formation in this context requires engineered T3 variants to support productive, state-dependent interdomain transfer, including efficient loading by A3 and compatible interactions with the flanking C-domains (acceptor C3 and donor C/E4).

We next tested whether top-performing designs from the SU2 prescreen translate to the full assembly line. We selected the highest-yielding AI-designed T domains across rounds 1–3 and model classes (AI-2; AI-12, AI-15, AI-27 and AI-31; and AI-38, AI-44 and AI-59) and installed each at the T3 position of GxpS-SZ17/18. We quantified production of **6** in 20-mL cultures in succinate XPP medium (Fig. 3a). All eight variants produced detectable **6**, indicating that *de novo* T-domain sequences can remain compatible with both upstream and downstream interfaces in the complete assembly line. Strikingly, two variants exceeded WT titers: AI-27 reached 257 % of WT (7.4 mg L^−1^) and AI-38 reached 144 % of WT (4.1 mg L^−1^). Both variants were functional in the SU2-only assay, yet their relative gains increased in the full-length system, indicating that prescreen performance in a reduced context does not fully predict behavior in the complete assembly line.

**Fig 3:**
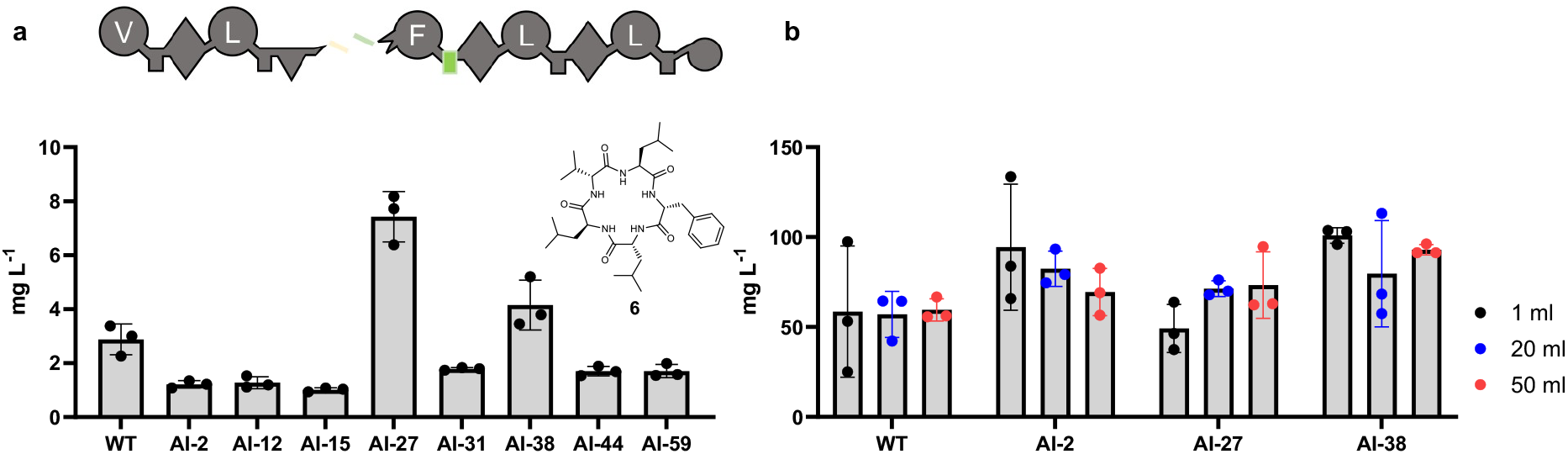
AI-designed T-domains support efficient production of 6 across variants and culture scales. **a**, Titer performance of AI-generated T-domain variants at the T3 position in the pentapeptide production system, measured in 20 mL cultures grown in succinate XPP medium. **b**, Scale-dependent performance of selected AI-designed T3 variants. Production titers of **6** were quantified at three culture volumes (1 mL, 20 mL, and 50 mL) in LB medium for WT and selected AI-designed variants (AI-2, AI-27, and AI-38). Titers (mg L^−1^) were determined by LC-MS/MS in SRM mode using an external calibration curve. Data represent mean ± SD from three independent biological replicates. Source data are provided with this paper.

Because the DBTL workflow uses titers of **6** as quantitative functional labels, we next assessed how cultivation conditions influence these measurements. Changing the culture medium substantially altered titers – in succinate XPP medium at 20 mL, WT production was ∼2.9 mg L^−1^ (Fig. 3a), whereas in LB at the same volume it increased to ∼50 mg L^−1^ (Fig. 3b). In LB, WT titers were largely insensitive to culture volume, remaining relatively stable from 1 to 50 mL. Across this range, AI-2 and AI-38 supported elevated titers at 1 mL with modest decreases at larger volumes, while remaining comparable to or slightly above WT at 50 mL. NRPSs carrying AI-27 tracked WT at small scale but increased modestly at 20–50 mL, reaching titers comparable to AI-2 at 50 mL. Together, these experiments demonstrate that absolute titers, and in some cases relative differences between variants, depend strongly on cultivation conditions. Accordingly, production titers should be interpreted as condition-dependent functional labels rather than context-independent measures of sequence performance. These findings highlight the need for standardized conditions to ensure internal consistency across DBTL rounds.

### AI-designed T-domains exhibit enhanced foldability and biochemical robustness

AI-2 emerged as a strong functional hit in the first design round in the SU2 assay and was the earliest design that exceeded WT activity; we therefore prioritized it for detailed biochemical and computational characterization.

The higher product formation supported by AI-2 relative to WT GxpS_T3 across multiple assay conditions (Fig. 2a; Fig. 3b) led us to examine whether generative design altered intrinsic developability and biochemical robustness of the T-domain. We therefore compared expression, solubility and folding behavior of AI-2 and WT GxpS_T3 across several domain contexts, including isolated T3/AI-2 and A3–T3/AI-2–C/E4-containing constructs with identical junction linkers. Across expression conditions, AI-2 consistently accumulated at higher levels and was readily recovered in the soluble fraction, whereas WT GxpS_T3 was predominantly insoluble and required denaturing purification (Fig. 4a–c).

**Fig 4:**
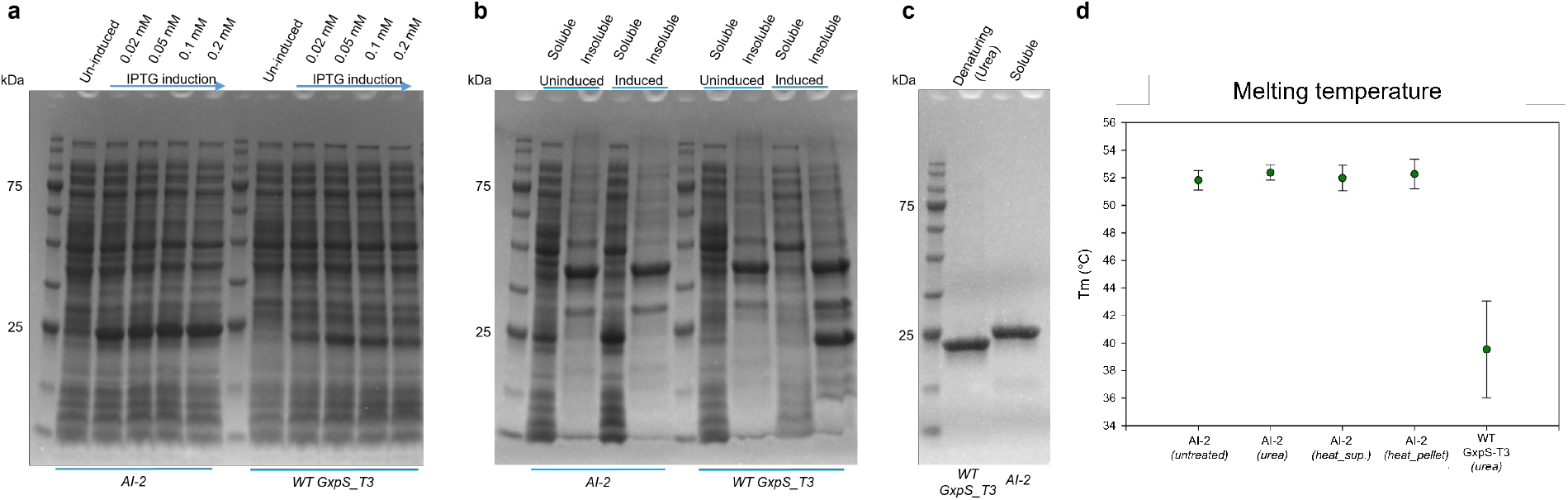
The AI-designed T-domain exhibits enhanced expression, solubility, and thermal stability compared to the wild-type domain. **a**, Expression levels of WT GxpS_T3 and AI-2 following induction in *E. coli*. **b**, Solubility analysis of WT GxpS_T3 and AI-2. Protein expression was induced with 0.05 mM IPTG and cultures were incubated at 18 °C for 16 h. Soluble (S) and insoluble (I) fractions are shown. **c**, Purification profiles of WT GxpS_T3 and AI-2. WT GxpS_T3 was purified under denaturing conditions due to insolubility, whereas AI-2 was recovered in soluble form. **d**, Thermal stability of WT GxpS_T3 and AI-2 was determined by thermal denaturation. Melting temperatures (Tm) were calculated as the midpoint of the unfolding transition. Data are shown as mean ± SD from three independent biological replicates, each representing the mean of three technical measurements.

After denaturation, WT GxpS_T3 largely aggregated upon refolding, eluting in the void volume by size-exclusion chromatography (SEC), and only a minor fraction recovered a soluble, native-like state (Supplementary Fig. S5a). By contrast, AI-2 refolded efficiently and recovered a native SEC profile (Supplementary Fig. S5b). Thermal denaturation further supported increased stability of AI-2 (Tm ∼52 °C) compared with refolded WT GxpS_T3 (Tm ∼40 °C) (Fig. 4d). Similar trends were observed for multi-domain constructs containing AI-2, which consistently showed improved expression and solubility relative to WT counterparts (Supplementary Fig. S6).

Together, these results identify AI-2 as a markedly more tractable T-domain scaffold than WT GxpS_T3, combining high soluble expression with efficient refolding and increased thermal stability. However, AI-2 did not maximize titers across all assembly-line contexts and cultivation conditions, indicating that improved fold robustness alone does not fully determine pathway output. Consistent with this separation between intrinsic developability and context-dependent productivity, epPCR diversification of AI-2 yielded a high fraction of improved variants, including the most active T-domain variant identified here (285% relative to WT; variant 123; Supplementary Data 1). This motivated a closer examination of how sequence changes reshape state-dependent interdomain contacts.

### Molecular dynamics simulations reveal state-dependent alterations of interdomain contacts in WT GxpS_T3 and AI-2

To probe catalytic-state-dependent dynamics at the T3 junction, we performed 100-ns atomistic molecular dynamics (MD) simulations of WT GxpS_T3 and AI-2 in three catalytic states capturing thiolation, condensation donor and condensation acceptor interfaces (Supplementary Figs. S7–S9), using homology models based on PDB 5T3D^53^, 9BE3^54^, and 8JBR^55^. Across states, both variants maintained comparable global structural stability of the A3-, T3- and C-domains, with similar backbone deviation and fluctuation profiles as well as stable measures of compactness and solvent exposure (Supplementary Figs. S7–S9). The T-domain core remained particularly stable (∼1.5–1.8 Å RMSD), whereas increased apparent motion arose primarily from flexible linker regions rather than destabilization of folded domains.

Despite similar global dynamics, contact analysis revealed clear, state-dependent differences in interdomain interaction organization (Fig. 5; Supplementary Table S3). In the thiolation state, WT GxpS_T3 engaged A3 through a distributed interface involving the C-terminal region of helix 1, a short helix within loop 1 and helix 2, yielding more frequent contacts (≥50 % occupancy) than AI-2, which adopted a more focused interaction dominated by helix 2 (Fig. 5a). In the condensation donor state, both variants docked at the C/E4 donor site in geometries consistent with catalysis, but AI-2 showed a higher contribution of polar and hydrogen-bond-mediated contacts, including interactions involving arginine residues on helices 2 and 3 (Fig. 5b). In the condensation acceptor state, interactions were dominated by helices 2 and 3 contacting the C3 acceptor site and A3 subdomain, with WT GxpS_T3 maintaining a higher number of persistent contacts (Fig. 5c). Complete residue-level distance and hydrogen-bond interaction tables for each state and replicate are provided in Supplementary Data 1.

**Fig 5:**
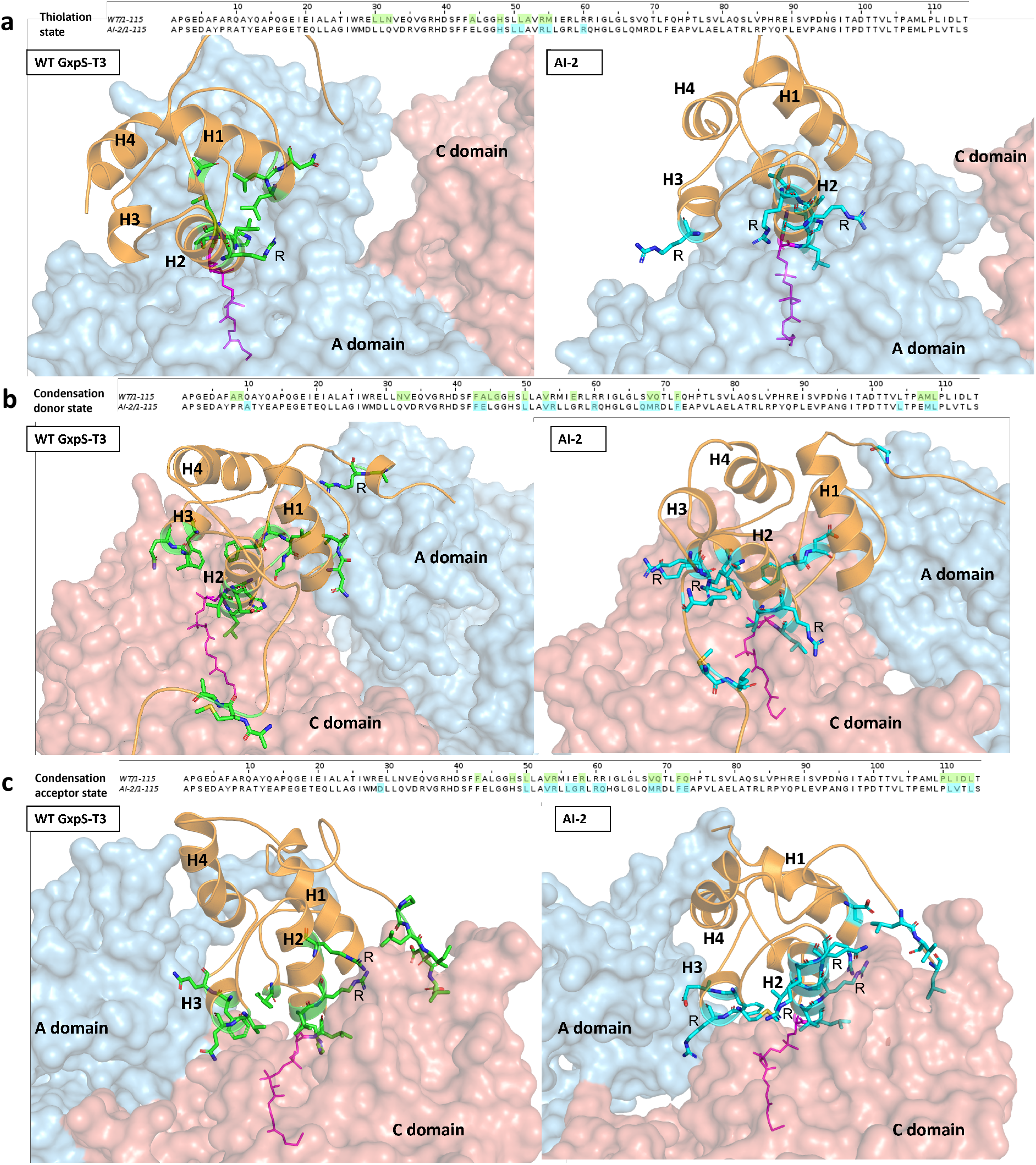
State-dependent interdomain interactions of WT GxpS_T3 and AI-2 T-domains across the NRPS catalytic cycle. Structural representations and sequence alignments showing interdomain contacts of WT and AI-2 T-domains in the thiolation **(a)**, condensation donor **(b)**, and condensation acceptor **(c)** states. Structural views are shown below (WT GxpS_T3 left, AI-2 right), with corresponding sequence alignments displayed above. Interacting residues identified by molecular dynamics simulations are shown as sticks and highlighted in green (WT GxpS_T3) and cyan (AI-2), and are marked in the sequence alignment using the same color scheme. Only R residues were labeled in the structural representations. The T-domain is shown in orange, the adenylation (A) domain in blue, the condensation (C) domain in salmon, and the PPant arm in magenta. Major structural elements of the T-domain are labeled H1–H4. Interactions were defined as hydrogen bonds with ≥30% occupancy and/or distance-based contacts with ≥80% occupancy across simulation trajectories.

Together, these simulations indicate that AI-2 preserves global fold stability while reshaping the distribution, persistence and chemical nature of interdomain contacts in a state-dependent manner. This observation is consistent with the context dependence observed *in vivo*, where the SU2-only system primarily samples A3 and C/E4 interfaces, whereas the full assembly line additionally requires coordinated engagement at the upstream C3 acceptor interface. These findings are consistent with recent mechanistic studies showing that NRPS carrier domains engage their partners through multiple weak and transient interactions that balance specificity with dynamic exchange ^56-58^.

### AI-designed T-domains show strong positional dependence within the GxpS assembly line

To test whether the state-dependent interface differences observed in MD translate into position-specific constraints *in vivo*, we selected three T-domain variants spanning design rounds and functional profiles: AI-2 as an early robust scaffold, AI-12 as a WT-like performer in the SU2 assay that underperformed in the full assembly line context, and AI-38 as a top-performing variant active in both assay configurations. Each variant was installed at positions T1, T2, T4, and T5 of GxpS to assess positional compatibility within the assembly line (Fig. 6). For each position, we quantified production of **6** relative to the native T-domain normally occupying that position (set to 100 %). At T1, AI-2 and AI-12 abolished detectable production of **6**, whereas AI-38 retained measurable activity (∼24 % of the native T1 control). At T2, AI-2 and AI-12 supported ∼42 % and ∼16 % of the native T2 control, respectively, while AI-38 was non-functional. By contrast, T4 tolerated substitution more broadly: AI-2 reached ∼43 % of the native T4 control, AI-12 exceeded the native control (∼137 %), and AI-38 produced ∼61 % of native T4 levels. T5 was the most permissive position, with AI-2 and AI-12 substantially surpassing the native T5 control (∼216 % and ∼272 %, respectively), whereas AI-38 again yielded no detectable production of **6** (Fig. 6).

**Fig 6:**
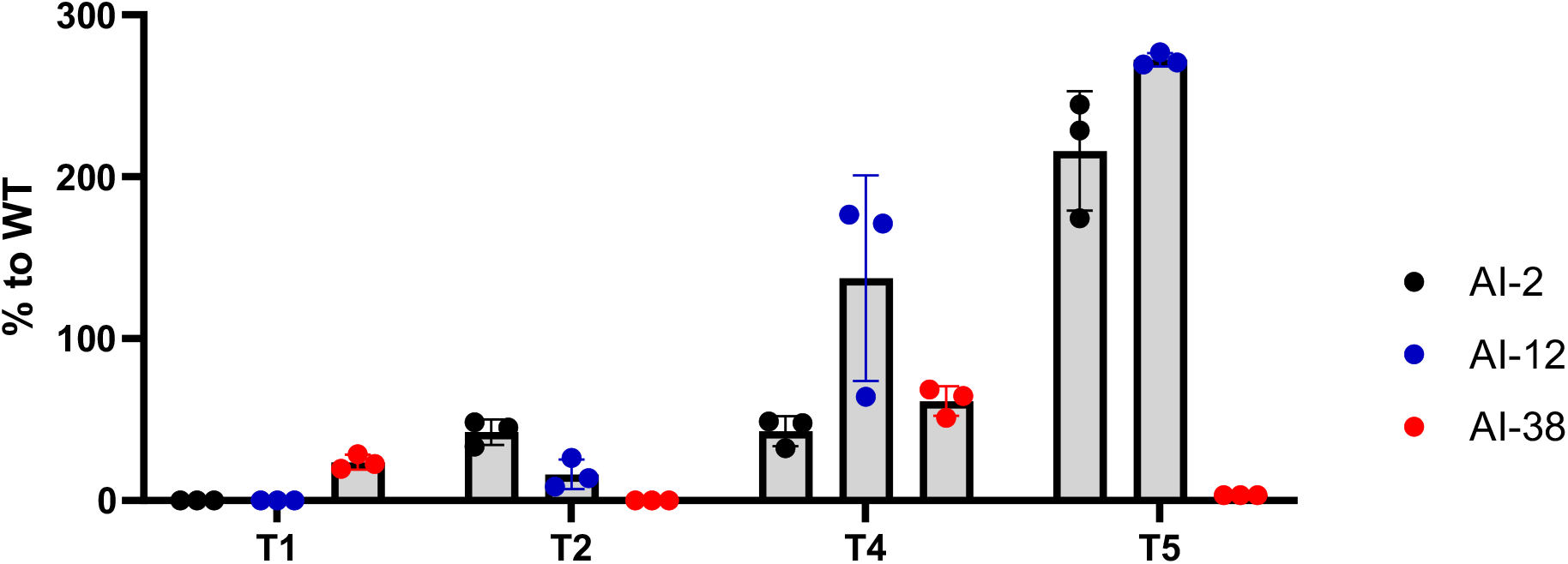
Production of 6 by GxpS constructs containing AI-designed T-domains at different positions. Titers are shown relative to the corresponding native T-domain at each position, which was set to 100 %. Bars show the mean ± SD from three independent biological replicates. Source data are provided with this paper.

This pronounced positional dependence is qualitatively consistent with T-domains encoding partner- and state-specific interface features, in agreement with the interdomain plasticity observed by MD in the T3 context (Fig. 5). Notably, our designs were generated in the T3 neighborhood, where the immediate downstream partner is the dual C/E4 domain, whereas at T2 the downstream partner is an ^L^C_L_-type C-domain (Fig. 1a). Given that ^L^C_L_- and dual C/E-domains differ in stereochemical requirements and catalytic mechanisms^59^, interfaces compatible with the T3–C/E4 junction may not transfer to the T2–C3 (^L^C_L_-type) junction. Consistent with this view, downstream-facing T-domain features co-vary with the identity and chemistry of downstream partner domains^39^. Together, these results indicate that T-domain compatibility in GxpS is position-dependent and strongly influenced by the identity of the immediate downstream partner.

### AI-designed T-domains can be leveraged to generate hybrid NRPSs

Having shown that the activity of AI-designed T-domains depends strongly on their local catalytic and interaction context within GxpS (Fig. 3a and Fig. 6), we next asked whether they can be used as practical engineering parts to support the construction of functional hybrid NRPSs. To this end, we generated chimeric assembly lines by recombining GxpS with modules derived from two distinct biosynthetic gene clusters - the xenotetrapeptide synthetase (XtpS) from *Xenohabdus nematophila* HGB081^60^ and the szentiamide synthetase (SzeS) from *Xenorhabdus szentirmaii* DSM 16338T^61^. These hybrid systems impose non-cognate A–T–C interfaces and therefore provide a stringent test of functional generalization, consistent with evidence that interdomain communication frequently limits NRPS engineering outcomes^57,62^.

In the first GxpS–XtpS hybrid NRPS assembly line (GxhS-1), the construct containing the native GxpS_T3 domain produced only minimal amounts of the tetrapeptide vLfV (**9**), corresponding to LC-MS peak areas of ∼2 × 10^4^. In contrast, GxhS-1 constructs carrying AI-2, AI-27, and AI-31 yielded markedly higher peak areas (>4.5 × 10^6^), whereas constructs carrying AI-12 and AI-32 showed no detectable peptide formation (Supplementary Fig. S11; Fig. 7a). The GxpS–SzeS hybrid (GshS), assembled using an XUT-based^39^ recombination strategy that fuses GxpS to the T4–TE region of SzeS, produced LC-MS peak areas of ∼4 × 10^6^ for the pentapeptide vLfYW (**16**), with the native Sze_T4 installed. Constructs carrying AI-2 and AI-12 increased peak areas to ∼5 × 10^6^ and ∼6.5 × 10^6^, respectively (∼25% and ∼65% increases relative to the native T-domain), while the construct carrying AI-31 performed comparably and those carrying AI-27 and AI-32 showed reduced product formation (<2 × 10^6^) (Supplementary Fig. S12; Fig. 7b).

**Fig 7:**
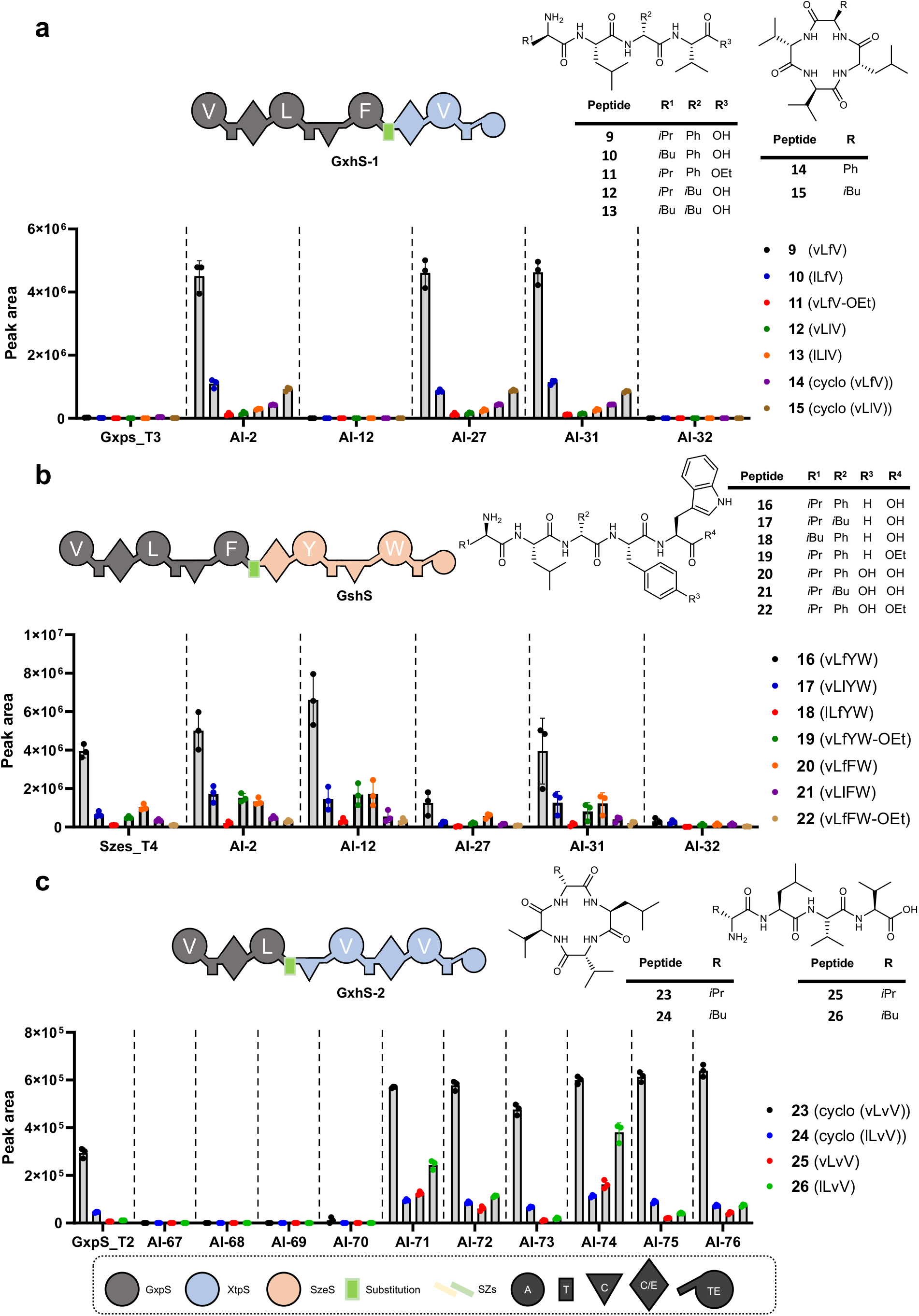
Production of peptides by hybrid NRPSs containing AI-designed T-domains. Bar plots show LC-MS peak areas corresponding to relative peptide abundance generated by hybrid NRPSs: **a**, GxhS-1 producing linear vLfV **(9)** as the dominant product; **b**, GshS producing vLfYW **(16)** as the dominant product; **c**, GxhS-2 producing cyclic vLvV **(23)** as the dominant product. Colored markers denote individual peptide species. Bars represent mean values from three independent biological replicates, with individual replicate measurements shown as overlaid points. LC-MS peak areas were used as a semi-quantitative measure of relative peptide abundance across matched experimental conditions. Source data are provided with this paper.

To test interface-specific design, T-domains optimized for ^L^C_L_-type condensation interfaces were evaluated in a second GxpS–XtpS hybrid (GxhS-2), in which GxpS_T2 was fused upstream of the C3-domain of XtpS to position the engineered carrier directly before an ^L^C_L_-type C-domain. In this context, NRPS constructs carrying the native GxpS_T2 supported production of cyclo(vLvV) (**23**), corresponding to LC-MS peak areas of ∼3 × 10^5^. Variants carrying AI-71 to AI-76 reached LC-MS peak areas of ∼6 × 10^5^, whereas constructs carrying AI-67 to AI-70 yielded no detectable product formation (Supplementary Fig. S13; Fig. 7c).

Because peak areas were compared under matched LC-MS conditions, these differences reflect relative changes in product formation. Notably, improved production was observed in a hybrid assembled via an XUT-style^39^ insertion architecture (Fig. 7b), showing that AI-designed carrier domains can outperform established evolution-guided NRPS engineering approaches.

Motivated by the pronounced context dependence observed across hybrid systems, we next performed a high-throughput evaluation of 76 AI-designed T-domains across multiple NRPS architectures, including GxhS-1, GxhS-2, and GshS and the native SzeS assembly line, which produces the pentapeptide cyclo(lTfvYW) (**27**) (szentiamide)^61^ (Supplementary Fig. S14a). All constructs were screened using a RapidFire LC-MS system, and production was assessed semi-quantitatively based on LC-MS peak areas corresponding to the expected main product (Supplementary Fig. S14b).

Across hybrid architectures, the identity of the downstream C-domain emerged as the primary determinant of compatibility (^L^C_L_-type C versus C/E). Constructs were classified as improved (production exceeding the corresponding WT control), inactive (no detectable product), or intermediate (detectable but ≤ WT levels). In constructs in which the transplanted T-domain was followed by a C/E-domain (GxhS-1 and GshS), many AI-designed T-domains increased production relative to the native reference (GxhS-1: 53 improved, 19 inactive; GshS: 15 improved, 9 inactive), whereas the set of designs tailored for ^L^C_L_-type C-domain interfaces (AI-67–AI-76) consistently yielded no detectable products. A similar pattern was observed in the native SzeS assembly line, which also contains a downstream C/E-domain: 4 constructs improved production relative to the native reference, 46 were inactive, and none of the AI-67– AI-76 constructs produced detectable peptide amounts. By contrast, in GxhS-2, where the transplanted T-domains precede an ^L^C_L_-type C-domain, only 5 constructs supported detectable production, including 3 of the ^L^C_L_-tailored designs, while most C/E-optimized T-domains failed.

To connect these functional trends to sequence space, we performed a sequence-distance-based phylogenetic analysis of T-domains annotated by downstream partner identity (Supplementary Fig. S10). AI-1–AI-66 clustered predominantly with WT T-domains associated with C/E interfaces, whereas AI-67–AI-76 grouped with T-domains linked to ^L^C_L_-type C-domains. This separation mirrors the functional patterns observed in the positional and hybrid experiments (Figs. 6 and 7) and suggests that the generative models capture downstream-partner-specific features present in natural T-domain sequences.

Together, these results show that AI-designed T-domains can support product formation in multiple non-cognate hybrid NRPS architectures, and that compatibility is primarily conditioned by the class of the immediate downstream C-domain (^L^C_L_-type versus C/E).

## Discussion

NRPSs remain among the most demanding enzyme systems to reprogram^17^. Even with detailed mechanistic and structural insight, predictable engineering is limited by the dynamic property that enables catalysis: productive turnover depends on transient, state-dependent interdomain interfaces^5^. Accordingly, the field has moved from motif-guided swaps toward structure-informed and evolution-guided recombination strategies that preserve compatible junctions^4,5,17,63,64^. Here we demonstrate that pretrained generative protein models, coupled to DBTL, add a practical capability: proposing new-to-nature carrier-domain sequences that are functional within defined catalytic neighborhoods.

Across 578 recombinant NRPS variants spanning minimal, full-length and hybrid architectures, *de novo* T-domain sequences frequently supported productive biosynthesis and, in defined contexts, improved titers relative to native carrier domains. In this work, we define “success” operationally as detectable product formation and, where applicable, increased production under matched assay conditions. Two principles emerged. First, sequence similarity to the native carrier domain had limited predictive value for function. Second, compatibility was governed by local junction context, with the identity of the immediate downstream partner domain emerging as a major determinant. This dependence was evident in positional transplantation within GxpS and in hybrid recombination experiments, where designs tailored for ^L^C_L_-type C-domain interfaces (AI-67–AI-76) failed in C/E contexts but supported activity when paired with an ^L^C_L_-type downstream partner. Together, these results indicate that junction identity is embedded in carrier-domain sequence space and should be treated as an explicit design variable.

Biochemical and computational analyses further reinforce an interface-centric interpretation. One of our earliest functional AI scaffolds (AI-2) displayed markedly improved developability, including higher soluble expression, efficient refolding and increased thermal stability, while MD simulations indicated preserved global fold stability but reorganization of state-dependent interdomain contact networks. Together, these results provide a mechanistic rationale for why a given carrier-domain sequence can be beneficial in one architecture yet neutral or detrimental in another. They argue for shifting from post hoc screening of hybrids to designing carrier domains for the intended local catalytic neighborhood as a route to context-conditioned NRPS engineering. More broadly, our results show that generative design can be applied to dynamic megasynth(et)ases by targeting the interface-encoding carrier domain.

Beyond enabling hybrid construction, generative carrier-domain design offers a potential route to improving pathway throughput. Because many biosynthetic gene clusters remain limited by low yields, increasing production can reduce material constraints that often bottleneck purification, structure elucidation and downstream bioactivity profiling^65^.

Our data also define a practical roadmap for making this capability increasingly predictive. Interface compatibility is inherently context dependent, so the goal is not universal interchangeability but reliable, junction-conditioned design. In addition, the quantitative labels used for DBTL are condition dependent: LC–MS readouts report pathway output but integrate interface effects with expression, growth physiology and cultivation conditions, and media or scale changes can shift absolute titers and, in some cases, variant rankings. This label variability can be viewed as “noise” from the perspective of learning a sequence-to-function mapping, motivating standardized assay conditions and larger, multi-condition datasets. Moving forward, deeper structural and kinetic interrogation of engineered junctions will be valuable, but complete structural coverage for all contexts may not be necessary if partner-aware predictors explicitly encode junction identity and are trained on expanding sequence– context–function datasets. Coupled to generative models, such predictors should progressively reduce reliance on iterative DBTL and enable increasingly prospective NRPS engineering, accelerating access to new-to-nature molecules and more efficient production of valuable natural products.

## Methods

### Bacterial Strain and growth conditions

All *E. coli* strains used in this study were given in Supplementary data 1. Strains were cultured in either liquid or solid LB medium (10 g L^−1^ tryptone, 5 g L^−1^ yeast extract, and 5 g L^−1^ NaCl, pH 6.8-7.2) and succinate XPP medium (36.9 g L^−1^ succinate, 20 mL M9 salt A, 20 mL M9 salt B, 20 g L^−1^ proteose peptone no. 3, 1 g L^−1^ Na-pyruvate, 0.2 % vitamin solution, and 0.1 % trace element solution). As a selection marker, kanamycin (50 µg mL^−1^), chloramphenicol (34 µg mL^−1^), ampicillin (50 µg mL^−1^), or sucrose (1 %) were added. Solid media were supplemented with 1 % (w/v) agarose.

Routine culturing was performed at 37 °C with 180 rpm shaking. For peptide production, promoter induction was conducted at 22 °C. The *E. coli* NEB Stable (NEB) was used for cloning, *E. coli* DH10B::*mtaA*^*66*^ for cloning and fermentation, and *E. coli* BL21 Star (DE3)(Invitrogen) for protein expression and purification.

### Plasmid construction and mutagenesis

All plasmids, constructs and primers were given in Supplementary data 1. Plasmids used as template DNA for construction of T-domain acceptor backbones (pMY-e4-0000-BsaI_noSZ-SacB, pET28a, pJW75_pACYC_ara_araE II_GxpS, pJW76_pCOLA_ara_tacI_II_GxpS, pJW76_T3_ESM3_2, pNA5_pCOLA_ara_tacI II_XCN1_2022_T2-TE, and pDC002) were propagated in *E. coli* and isolated using the Monarch^®^ Plasmid Miniprep Kit (NEB).

DNA fragments were amplified using Q5^®^ Hot Start High-Fidelity DNA Polymerase (NEB) or 2× Phanta Max Master Mix (Vazyme). PCR reactions were treated with DpnI (NEB) to remove methylated template DNA. PCR products were verified by electrophoresis on 1 % agarose gels. DNA was purified using the Monarch^®^ DNA Gel Extraction Kit (NEB) or Monarch^®^ PCR & DNA Cleanup Kit (NEB).

Purified fragments were assembled with NEBuilder^®^ HiFi DNA Assembly Master Mix (NEB) to generate acceptor plasmids containing BsaI sites for subsequent Golden Gate cloning.

AI-designed T-domain sequences were obtained by commercial gene synthesis (Eurofins Genomics or GenScript Biotech) and inserted into *BsaI-*flanked acceptor plasmids by Golden Gate assembly using BsaI (NEB) and T4 DNA ligase (Promega). Golden Gate reactions were performed with 80 cycles of 37 °C for 2 min and 16 °C for 5 min, followed by 50 °C for 5 min and 80 °C for 10 min, and then cooled to 12 °C. Negative selection of non-recombined clones was performed via the *sacB* cassette on media containing 1 % sucrose. Assembled plasmids were transformed by heat shock into *E. coli* DH10B::*mtaA, E. coli* BL21 Star (DE3), or NEB Stable cells, as indicated for each experiment.

Error-prone mutagenesis libraries based on WT GxpS_T3, AI-2, AI-12, and AI-15 were generated using the GeneMorph II Random Mutagenesis Kit (Agilent) according to the manufacturer’s instructions, targeting mutation frequencies of 0-4.5 or 4.5-9 mutations per kb.

### Fermentation and extraction

For peptide production, cultures were prepared either in 96-deep-well plates (**1** and **2**) screening) or in shake flasks for **6** and hybrid NRPS-derived peptides (**9-26**), as described below. Unless otherwise stated, cultures were harvested by centrifugation at 4 °C (4000 rpm, 10 min) to pellet cells (including XAD-16 resin where applicable), extracted with methanol at a 1:1 (v/v) ratio relative to culture volume, and clarified by centrifugation prior to LC-MS/MS analysis.

### 1 and 2 screening

96-deep-well plates were filled with 1 mL succinate XPP medium per well supplemented with 50 µg mL^−1^ kanamycin and 2 mM L-arabinose, inoculated with 20 µL overnight culture, and incubated at 22 °C with shaking at 1100 rpm for 48 h. Pellets were extracted by shaking at 15 °C and 1100 rpm for 40 min, and clarified by centrifugation at 4 °C (4000 rpm, for 20 min).

### Production of 6 and hybrid NRPSs-derived peptides (9-26)

For production of **6**, cultures were grown in either 20 mL succinate XPP medium or in 1 mL, 20 mL, and 50 mL LB medium, each supplemented with 34 µg mL^−1^ chloramphenicol and 50 µg mL^−1^ kanamycin, 2mM L-arabinose and 2 % (v/v) XAD-16 resin. For production of hybrid NRPS-derived peptides (**9-26**), cultures were grown in 20 mL succinate XPP medium supplemented with 2 mM L-arabinose and 2% (v/v) XAD-16 resin. Chloramphenicol (34 µg mL^−1^) was used for compounds **9-15** and **23-26**, whereas kanamycin (50 µg mL^−1^) was used for compounds **16-22**. Cultures were inoculated with 2 % (v/v) overnight culture and incubated at 22 °C and 180 rpm for 48 h. Pellets were extracted by shaking at 15 °C and 250 rpm for 40 min and clarified by centrifugation at 4 °C (15000 rpm, 20 min) prior to analysis.

### Synthesis of reference peptide synthesis and purification

**6** was synthesized using standard Fmoc solid-phase peptide synthesis (SPPS) on 2-chlorotrityl chloride resin, followed by cleavage, solution-phase macrocyclization. The product was purified by reverse-phase HPLC. Peptide identity and purity were confirmed by high-resolution mass spectrometry. Detailed synthesis, purification, and analytical procedures are provided in the Supplementary Methods.

### Targeted LC-MS/MS quantification

Tripeptides **1** and **2** and pentapeptide **6** were quantified on a Vanquish Flex UHPLC system coupled to a TSQ Altis Plus triple-quadrupole mass spectrometer (Thermo Fisher Scientific) operated in SRM mode. Separations were performed on a Hypersil GOLD C18 column (1.9 μm, 100 × 2.1 mm; Thermo Fisher Scientific) at 40 °C with a flow rate of 0.700 mL min^−1^ using solvent A (water supplemented with 0.1 % formic acid) and solvent B (acetonitrile supplemented with 0.1 % formic acid) The gradient for tripeptides **1** and **2** was 10 % B (0-0.2 min), 10-90 % B (0.2-0.4 min), 90 % B (0.4-1.2 min), and 10 % B (1.2-1.6 min). The gradient for pentapeptide **6** was 10 % B (0-0.2 min), 10-90 % B (0.2-0.3 min), 90 % B (0.3-1.4 min), and 10 % B (1.4-1.8 min). MS conditions were as follows: vaporizer temperature 350 °C, ion transfer tube temperature 380 °C, sheath gas 30, auxiliary gas 10, sweep gas 2. Mass transitions and further parameters are given in Supplementary Table S1. The amount of peptide was analyzed using a calibration curve ranging from 0.016 to 4 µM. For pentapeptide **6**, signals were normalized against the internal standard diphenhydramine (10 nM).

### LC-MS/MS analysis

Other peptide products (**9-26**) were analyzed on a Vanquish UHPLC system (Thermo Fisher Scientific) coupled to an ESI-QqTOF impact II mass spectrometer (Bruker Daltonics) operated in auto MS/MS mode.

Separations were performed using an ACQUITY UPLC BEH C18 VanGuard pre-column (130 Å, 1.7 µm, 5 × 2.1 mm; Waters), an ACQUITY UPLC BEH C18 analytical column (130 Å, 1.7 µm, 100 × 2.1 mm; Waters), and a KrudKatcher Ultra in-line filter (2.0 µm depth filter × 0.004 in ID; Waters) at 45 °C with a flow rate of 0.600 mL min^−1^. Solvent A was water with 0.1 % formic acid and solvent B acetonitrile with 0.1 % formic acid. The gradient was 5 % B (0–0.5 min), 5–95 % B (0.5–18.5 min), 95 % B (18.5–20.5 min), 95–5 % B (20.5–20.8 min), and 5 % B (20.8–22.5 min). Peak areas were integrated from extracted-ion chromatograms (EICs) and used for relative comparison across samples (no absolute quantification).

### MS1 settings

Mass spectra were acquired in centroid and profile modes over an m/z range of 50–1300 at 12 Hz. Source parameters were: end plate offset 500 V, capillary voltage 4500 V, nebulizer gas pressure 0.4 bar, dry gas flow 4 L min^−1^, and dry temperature 180 °C. Ion transfer and quadrupole settings were: funnel 1 RF 400 Vpp, funnel 2 RF 600 Vpp, ion energy 4 eV, and low-mass cut 150 m/z.

### MS/MS settings

MS/MS spectra were acquired at 2 Hz. The precursor list targeted the masses (Supplementary table S1) and was evaluated every 0.5 s, with a minimum precursor intensity threshold of 2500 counts. Collision-induced dissociation (CID) energies were set to a basic value of 50 and tune collision energy set to star 7 eV through end 200 eV.

### RapidFire LC-MS screening

Samples were prepared in a sealed 96-well plates and immediately prior to analysis centrifuged (3000 g, 15 min, 16°C). Samples were analyzed by high-throughput mass spectrometry using an Agilent RapidFire system (G9532A; Agilient) coupled to a TOF mass spectrometer equipped with an ESI (G6230B Dual AJS ESI; Agilient). The RapidFire platform aspirated samples directly from 96-well plates and a cleanup using solid-phase extraction (SPE, Agilent C4 Cartridge type A) was performed prior to elution into the MS. Each sample was aspirated from the well (aspiration time: 600 ms) and loaded onto the C4 cartridge using an aqueous load/wash solvent (Solvent A: water with 0.1 % formic acid). The cartridge was washed (wash/load time: 3000 ms), followed by elution (elution time: 6000 ms) of retained analytes into the TOF-MS using an organic elution solvent (Solvent B: acetonitrile with 0.1 % formic acid). The cartridge was then re-equilibrated before the next injection (re-equilibration time: 500 ms). TOF-MS data were acquired in positive and negative ion mode over an m/z range of 100-3000 m/z with a rate of 3 spectra s^−1^. Source conditions were set to: gas temperature 325 °C, drying gas flow 8 L min^−1^, nebulizer 35 psi, capillary voltage 1750 V and Nozzle voltage 1200 V. Mass calibration/reference mass correction was performed using Agilent ESI-L LC-MS Tuning Solution. Data was processed in Agilent RapidFire software and analyzed with MZmine software version 4.0.3^67^.

### Biochemical characterization of AI-2 and WT GxpS_T3 variants

Recombinant A3-T3-C/E4, A3-T3, T3-C/E4 and isolated T3 constructs corresponding to WT GxpS_T3 and AI-2 variants were cloned into pET28a-derived vectors encoding an N-terminal His_6_ tag followed by a TEV protease site. Proteins were expressed in *E. coli* BL21 Star (DE3) and purified by Ni–NTA affinity chromatography followed by size-exclusion chromatography. WT GxpS_T3 was purified under denaturing conditions and refolded by either rapid SEC-mediated urea removal or stepwise dialysis.

Protein solubility was assessed by SDS–PAGE analysis of soluble and insoluble fractions. Thermal stability was determined by nanoDSF, and melting temperatures (T_m_) were calculated from the first derivative of the fluorescence ratio (350/330 nm). Detailed cloning procedures, expression and solubility protocols, purification and refolding protocols, thermal stability measurements, and primer sequences are provided in the Supplementary Methods.

### Generative Design and Computational Prioritization of T-domains

*De novo* T-domain sequences were generated using three pretrained generative protein models: ESM3^11^, and EvoDiff^12^, ProteinMPNN^13^. In all cases, a conditional design strategy was applied in which the flanking A and C domains of the A-T-C tridomain were kept fixed and provided as structural and/or sequence context, while the T-domain region was masked and regenerated. This ensured that candidate sequences were explicitly conditioned on their intended architectural neighborhood. Generated sequences were evaluated *in silico* using multiple criteria, including sequence identity to WT GxpS_T3, structural plausibility assessed by ESMFold^52^ (mean pLDDT and TM-score), and perplexity under the ESM2 650M^52^ protein language model. These metrics were used to assess foldability, structural similarity, and biological plausibility prior to experimental testing.

To guide candidate prioritization in Round 3, we implemented surrogate-model–based active learning using experimentally measured sequence–activity data from Rounds 1 and 2 and from error-prone mutagenesis libraries. Labeled T-domain sequences evaluated in the SU2-only assay were used for training. Multiple modeling strategies were benchmarked, including linear regression on pretrained protein language model embeddings, zero-shot scoring approaches, and fine-tuning of ESMC^68^ using regression and contrastive objectives. Model performance was evaluated based on ranking quality (Spearman correlation), and two complementary strategies, linear regression on ESMC embeddings and contrastive fine-tuning, were selected for candidate prioritization.

For Round 3, new sequences were generated using ESM3 and EvoDiff, filtered using ensemble surrogate predictions and uncertainty estimates, and ranked according to predicted fitness. Top-ranked candidates were selected for experimental validation.

Detailed model architectures, training procedures, hyperparameters, benchmarking results, and filtering criteria are provided in Supplementary Notes.

### Homology Modeling of NRPS ATC tri-domain modules

Homology models of the ATC tri-domain module of the GxpS nonribosomal peptide synthetase were generated for both the WT GxpS_T3 and the AI-2 variants using MODELLER^69^. Modeling was performed to preserve the native multi-domain modular context of the NRPS, enabling explicit representation of interdomain interfaces that govern catalytic function.

To capture catalytically relevant conformations, separate ATC models were constructed for three experimentally characterized states of the NRPS catalytic cycle: the thiolation (loading) state, the condensation donor state, and the condensation acceptor state. Experimentally determined NRPS structures corresponding to these states were used as templates: the holo-EntF structure in the thioester-forming conformation (PDB 5T3D^53^) for the thiolation state, the pre-condensation donor state of LgrA (PDB 9BE3^54^) for the condensation donor state, and the condensation acceptor state of McyA2 (PDB 8JBR^55^).

Sequence alignments were generated using MOE^70^, focusing on conserved regions of the adenylation, thiolation, and condensation domains, with particular emphasis on the T-domain core and its flanking linker regions. Pairwise sequence identity between the GxpS T-domain core and the corresponding regions of the template structures ranged from approximately 22-35 %, which is typical for NRPS T-domain modeling. Conserved catalytic and structural motifs, including the phosphopantetheine-accepting serine of the T-domain, were preserved across all models.

In all homology models, the thiolation domain was represented in its holo form containing the PPant arm, without a covalently attached amino acid or peptide intermediate. Accordingly, all models represent pre-organized catalytic states prior to substrate transfer or condensation, rather than chemically reacting intermediates.

For each catalytic state, multiple ATC models were generated, and representative models were selected based on MODELLER statistical scores and structural consistency of interdomain interfaces. These ATC models served as starting structures for molecular dynamics simulations.

### Molecular Dynamics (MD) Simulations

All MD simulations were performed using AMBER24^71^. The ATC module models were parameterized with the ff19SB^72^ force field and solvated in a truncated octahedral box of OPC^73^ water with a minimum buffer distance of 10 Å from the protein surface. Systems were neutralized by the addition of counterions.

Energy minimization was carried out in two stages. First restraining the protein while minimizing solvent and ions, followed by unrestrained minimization of the entire system. Systems were then gradually heated from 0 to 300 K over 100 ps under constant-volume conditions with positional restraints applied to the protein. This was followed by density equilibration and a 5 ns equilibration phase under NPT conditions at 300 K and 1 atm, during which restraints were removed.

Production MD simulations were subsequently performed for 100 ns for each ATC model and catalytic state under NPT conditions using a 2 fs timestep. All covalent bonds involving hydrogen atoms were constrained using the SHAKE algorithm. Long-range electrostatic interactions were treated using the particle mesh Ewald method, and a cutoff of 10 Å was applied for nonbonded interactions. Temperature was maintained using a Langevin thermostat, and pressure was controlled using isotropic pressure coupling.

Because the starting structures were derived from homology modeling and represent pre-organized catalytic states within an ATC tri-domain module, the MD simulations were used to compare relative conformational stability, dynamic behavior, and interdomain interaction trends between ATC constructs containing either WT GxpS_T3 or AI-2, rather than to infer absolute structural accuracy or model chemical reaction steps.

Trajectory analyses were performed using cpptraj^74^ from the AMBER software suite. Backbone structural stability was assessed using RMSD, and residue-level flexibility was evaluated using RMSF. Global structural compactness and solvent exposure were quantified using Rg and SASA, respectively. Interdomain interactions were characterized by hydrogen bond analysis using geometric criteria (donor–acceptor distance ≤3.0 Å and angle ≥135°), and by distance-based residue–residue contact analysis, defined as residue pairs containing at least one heavy-atom pair within 4.5 Å for a defined fraction of the simulation trajectory.

### Phylogenetic reconstruction

Maximum-likelihood phylogenies were inferred using IQ-TREE v2.0.7 (multicore). Tree 1 was constructed from the full-length T-domain amino-acid multiple sequence alignment (129 sequences; 189 aligned positions), and Tree 2 from the second-half T-domain alignment. Sequence identifiers and amino-acid sequences used for phylogenetic analyses are provided in Supplementary data 1. Model selection was performed with ModelFinder Plus (-m MFP), and the best-fit substitution model was LG+F+R6 (chosen according to BIC). Branch support was assessed using 1,000 ultrafast bootstrap replicates (UFBoot2; -bb 1000) and 1,000 SH-aLRT replicates (-alrt 1000). Analyses were run in amino-acid mode (-st AA) with automatic thread allocation (-nt AUTO; seed 368473). Trees were visualized and annotated using iTOL, including color-coding by downstream domain type.

## Data availability

All plasmids, primers, strains, construct sequences and peptide lists are provided in Supplementary Data 1. Source data underlying the main figures are provided with this paper. Molecular dynamics simulation trajectories, associated input files and raw LC–MS data are available from the corresponding authors upon reasonable request.

## Code availability

Custom code was used for generative design, surrogate modeling, candidate prioritization and downstream analyses, as described in the Supplementary Information. The code is available from the corresponding author upon reasonable request. All third-party software and relevant parameters are detailed in the Methods and Supplementary Information.

## Acknowledgments

Andreas Keller (ZBI) for reviewing the manuscript and valuable feedback, Stefan Bauer (MCML) for fruitful discussions, Angela Sester (HIPS) for LC-MS support, and David Schwegler (Myria Biosciencs AG) for Lab support. This study was supported by the Helmholtz Gemeinschaft Deutscher Forschungszentren (HGF) by funding the Helmholtz Young Investigators Group of Kenan Bozhüyük [VH-NG-19-30] and the Innosuisse Innovation Project AIMMS 109.305 IP-LS.

## Contributions

K.A.J.B. conceived the study. O.V.K., D.K. and K.A.J.B. supervised the computational and experimental work. D.K. supported computational analyses. K.G. and V.P. performed sequence generation and surrogate modeling. E.F.B. performed molecular dynamics simulations and analysis. E.F.B., S.B., S.C., R.M. and H.A.M. performed NRPS engineering, cloning and in vivo testing of T-domain variants. S.B., R.H., S.S. and A.M.K. carried out LC– MS analytics and quantitative data analysis. D.W.H. and P.R. performed biochemical characterization of T-domain constructs. A.K.H.H and W.A.M.E. supervised GameXPeptide synthesis and analyzed the corresponding data; G.B. synthesized GameXPeptide. E.F.B., S.B., K.G. and K.A.J.B. wrote the manuscript with input from all authors.

