## Supplementary Information for "Generative AI designs functional thiolation domains for reprogramming non-ribosomal peptide synthetases"

1 Helmholtz Institute for Pharmaceutical Research Saarland (HIPS), Helmholtz Centre for Infection Research (HZI), PharmaScienceHub (PSH), Campus E8.1, 66123, Saarbrücken, Germany
2 Myria Biosciences AG, Tech Park Basel, Hochbergstrasse 60C, 4057, Basel, Switzerland
3 Center for Bioinformatics, Saarland University,Informatics Campus E2.1, 66123, Saarbrücken, Germany
4 Medical Faculty, Saarland University, Gebäude 74, 66421, Homburg, Germany
5 Department of Molecular Structural Biology, Helmholtz Centre for Infection Research, Inhoffenstr.7, 38124, Braunschweig, Germany
6 Technical University of Munich - Helmholtz AI - Munich Center for Machine Learning (MCML), Friedrich-Ludwig-Bauer-Str. 5, 85748, Garching b. München, Germany
7 Deutsches Zentrum für Infektionsforschung (DZIF), Inhoffenstr. 7, 38124, Braunschweig, Germany
8 Saarland University, Department of Pharmacy, Campus E8.1, 66123, Saarbrücken, Germany
9 Laboratory of Bioprocessing, D-BSSE, Eidgenössische Technische Hochschule (ETH) Zürich, Klingelbergstrasse 48, 4056, Basel, Switzerland
10 Spoken Language Systems (LSV), Saarland University, Campus C7.1, 66123, Saarbrücken, Germany
† Co-first authors
* Corresponding author

### Supplementary Information

#### Supplementary Tables

**Supplementary Table S1**. ESI-MS data of biosynthesized peptides, including peptide codes, calculated m/z values, molecular formulas, and literature references[1-4]**.** Construct abbreviations are defined as follows: GxpS SU2, GameXPeptide synthetase (GxpS) subunit 2 expressed alone; GxpS full, co-expression of GxpS subunits 1 and 2; GxhS-1 and GxhS-2, hybrid synthetases assembled from GxpS and xenotetrapeptide synthetase (XtpS) modules (variant 1 or 2, respectively); and GshS, a hybrid synthetase assembled from GxpS and szentiamide synthetase (SzeS) modules.

| NRPS system | Peptide | MS calculated $[M+H]^+$ | Molecular formula | Compound | Reference |
|---|---|---|---|---|---|
| GxpS SU2 | **1, 2** | 392.2544 | $C_{18}H_{35}N_3O_4$ | fIL/FIL | [1] |
| | **3, 4** | 358.2700 | $C_{21}H_{33}N_3O_4$ | lIL/LIL | [1] |
| GxpS full | **5** | 552.4119 | $C_{29}H_{53}N_5O_5$ | cyclo(vLlIL) | [2] |
| | **6** | 586.3963 | $C_{32}H_{51}N_5O_5$ | cyclo(vLfIL) | [2] |
| | **7** | 566.4276 | $C_{30}H_{55}N_5O_5$ | cyclo(lLlIL) | [2] |
| | **8** | 600.4119 | $C_{33}H_{53}N_5O_5$ | cyclo(lLfIL) | [2] |
| GxhS-1 | **9** | 477.3071 | $C_{25}H_{40}N_4O_5$ | vLfV | This study |
| | **10** | 491.3228 | $C_{26}H_{42}N_4O_5$ | lLfV | This study |
| | **11** | 505.3384 | $C_{27}H_{44}N_4O_5$ | vLfV-OEt | This study |
| | **12** | 443.3228 | $C_{22}H_{42}N_4O_5$ | vLlV | This study |
| | **13** | 457.3384 | $C_{23}H_{44}N_4O_5$ | lLlV | This study |
| | **14** | 459.2966 | $C_{25}H_{38}N_4O_4$ | cyclo(vLfV) | [3] |
| | **15** | 425.3122 | $C_{22}H_{40}N_4O_4$ | cyclo(vLlV) | [3] |
| GshS | **16** | 727.3814 | $C_{40}H_{50}N_6O_7$ | vLfYW | This study |
| | **17** | 693.3970 | $C_{37}H_{52}N_6O_7$ | vLlYW | This study |
| | **18** | 741.3970 | $C_{41}H_{52}N_6O_7$ | lLfYW | This study |
| | **19** | 755.4127 | $C_{42}H_{54}N_6O_7$ | vLfYW-OEt | This study |
| | **20** | 711.3865 | $C_{40}H_{50}N_6O_6$ | vLfFW | This study |
| | **21** | 677.4021 | $C_{37}H_{52}N_6O_6$ | vLlFW | This study |
| | **22** | 739.4178 | $C_{42}H_{54}N_6O_6$ | vLfFW-OEt | This study |
| GxhS-2 | **23** | 411.2966 | $C_{21}H_{38}N_4O_4$ | cyclo(vLvV) | [3] |
| | **24** | 425.3122 | $C_{22}H_{40}N_4O_4$ | cyclo(lLvV) | [3] |
| | **25** | 429.3071 | $C_{21}H_{40}N_4O_5$ | vLvV | This study |
| | **26** | 443.3228 | $C_{22}H_{42}N_4O_5$ | lLvV | This study |
| SzeS | **27** | 838.4134 | $C_{45}H_{55}N_7O_9$ | lTfvYW | [4] |

**Supplementary Table S2.** LC-ESI-MS/MS conditions and parameters for reaction monitoring of **1,2, 6**, and diphenhydramine (**DPH**).

| Analyte | Q1 mass [Da] | Q3 mass [Da] | Collision Energy [V] | Tube lens offset [V] | Spray Voltage [V] | Retention Time [min] |
|---|---|---|---|---|---|---|
| **1 and 2** | 390.217 | 129.967 | 21.9 | 92 | 3700 | 1.08 and 1.1 |
| | | 259.167 | 18.7 | | | |
| **6** | 586.367 | 261.133 | 21.9 | 79 | 4000 | 1.35 |
| | | 374.217 | 18.7 | | | |
| **DPH** | 256.05 | 164.967 | 41.8 | 30 | 4000 | 1.05 |
| | | 166.967 | 13.0 | | | |

**Supplementary Table S3**. Quantification of interface contact persistence across catalytic states. Residue–residue contact statistics derived from MD simulations of the thiolation, condensation donor (C_donor), and condensation acceptor (C_acceptor) states for WT GxpS_T3 and AI-2 constructs (two independent replicates each). For each interface (A–T or T–C), "≥50% occupancy" denotes the number of residue–residue contact pairs present in ≥50 % of analyzed MD frames, while "≥80% occupancy" indicates the number of highly persistent contacts present in ≥80 % of frames. Contacts were defined using a heavy-atom distance cutoff of 4.5 Å and evaluated every 10th frame over 100 ns trajectories (1,000 frames analyzed per replicate). Interfaces are defined as follows: **A–T** denotes the interface between the A-domain and the T-domain, whereas **T–C** denotes the interface between the T-domain and the C-domain.

| State | Interface | Construct | ≥50% occupancy | ≥80% occupancy |
|---|---|---|---|---|
| Thiolation_Rep1 | AT | WT | 28 | 19 |
| Thiolation_Rep2 | AT | WT | 29 | 16 |
| Thiolation_Rep1 | AT | AI-2 | 26 | 10 |
| Thiolation_Rep2 | AT | AI-2 | 20 | 12 |
| C_donor_Rep1 | AT | WT | 10 | 6 |
| C_donor_Rep2 | AT | WT | 10 | 5 |
| C_donor_Rep1 | TC | WT | 47 | 25 |
| C_donor_Rep2 | TC | WT | 38 | 21 |
| C_donor_Rep1 | AT | AI-2 | 2 | 1 |
| C_donor_Rep2 | AT | AI-2 | 13 | 6 |
| C_donor_Rep1 | TC | AI-2 | 43 | 23 |
| C_donor_Rep2 | TC | AI-2 | 30 | 17 |
| C_acceptor_Rep1 | AT | WT | 16 | 8 |
| C_acceptor_Rep2 | AT | WT | 6 | 2 |
| C_acceptor_Rep1 | TC | WT | 38 | 24 |
| C_acceptor_Rep2 | TC | WT | 52 | 31 |
| C_acceptor_Rep1 | AT | AI-2 | 4 | 2 |
| C_acceptor_Rep2 | AT | AI-2 | 4 | 1 |
| C_acceptor_Rep1 | TC | AI-2 | 28 | 17 |
| C_acceptor_Rep2 | TC | AI-2 | 43 | 33 |

#### Supplementary Figures

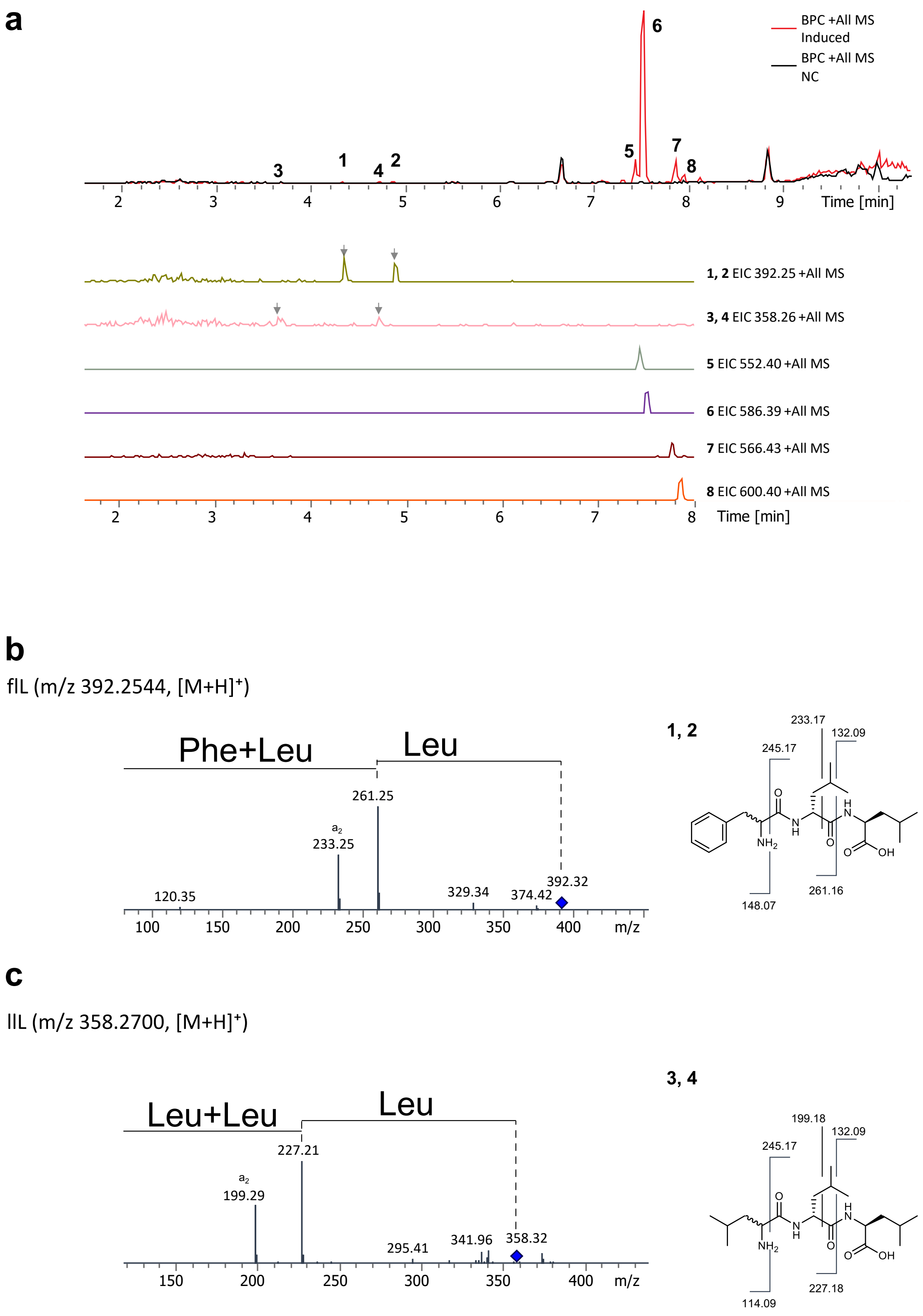

**d**

vLlIL (m/z 552.4119, [M+H]$^+$)

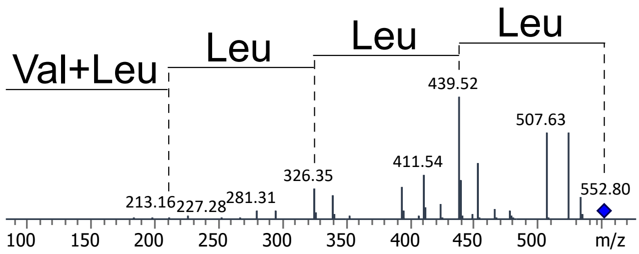

**5**

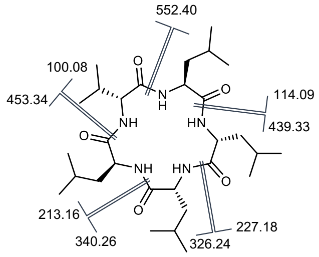

**e**

vLfIL (m/z 586.3963, [M+H]$^+$)

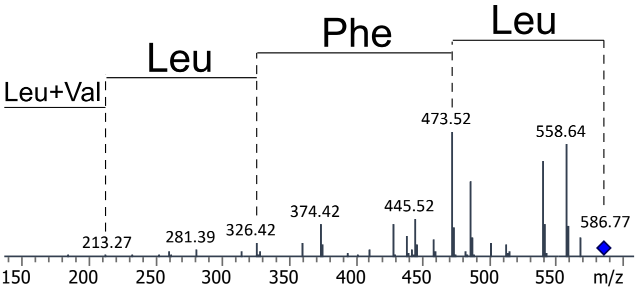

**6**

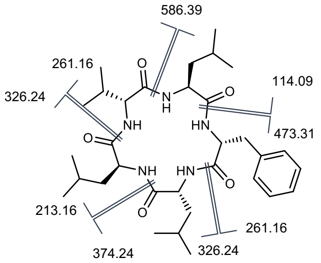

**f**

ILlIL (m/z 566.4276, [M+H]$^+$)

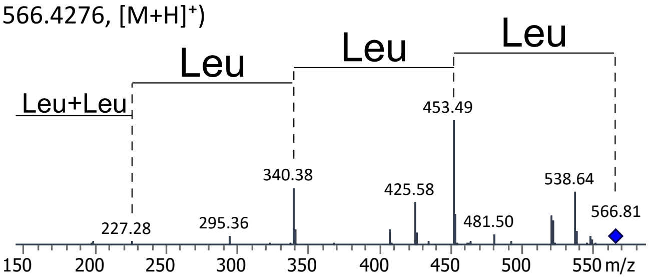

**7**

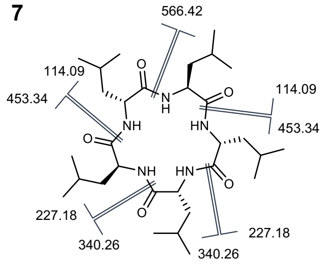

**g**

ILfIL (m/z 600.4119, [M+H]$^+$)

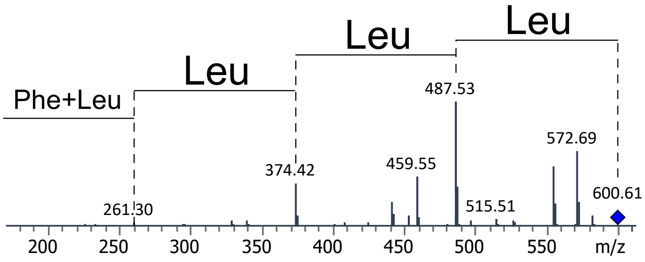

**8**

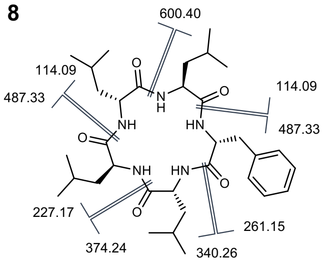

**Supplementary Figure S1**. Base peak chromatograms (BPC) and extracted ion chromatograms (EIC) of Type S GxpS expressed peptides. **a,** Base peak chromatograms (BPCs) and extracted ion chromatograms (EICs; extracted at the expected m/z values) for peptides produced by GxpS expression. Distinct chromatographic peaks corresponding to individual peptide products are labeled **1–8**. **b–g** Tandem mass spectrometry (MS/MS) spectra of peptides **1–8** with annotated fragment ions, confirming product identity.

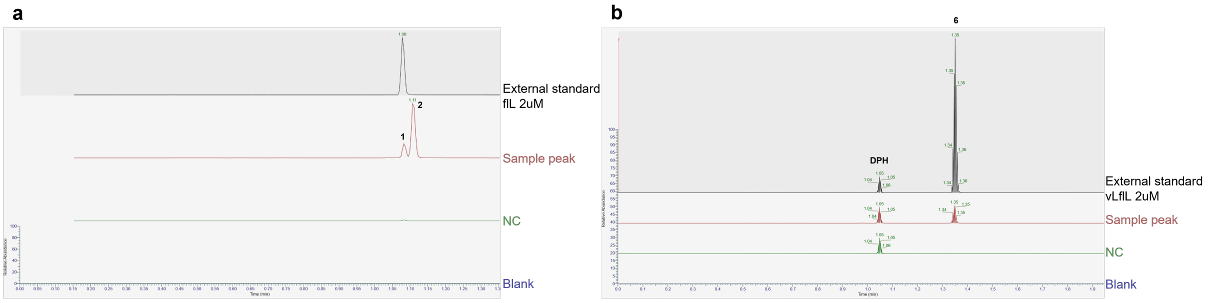

**Supplementary Figure S2**. LC-ESI-MS/MS chromatograms comparing producer extracts to external standards. **a**, ECIs for tripeptides **1** and **2**. The SU-2 only construct produced two major peaks corresponding to a mixture of linear tripeptides products[1]. **b,** EICs for pentapeptide **6.** Traces show the 2mM external standard (black), producer sample (red), negative control containing diphenhydramine (DPH; in **b**), and blank (blue). Distinct epimers were distinguished based on their retention times[1]. LC-ESI-MS/MS conditions and parameters are listed in Supplementary table S2.

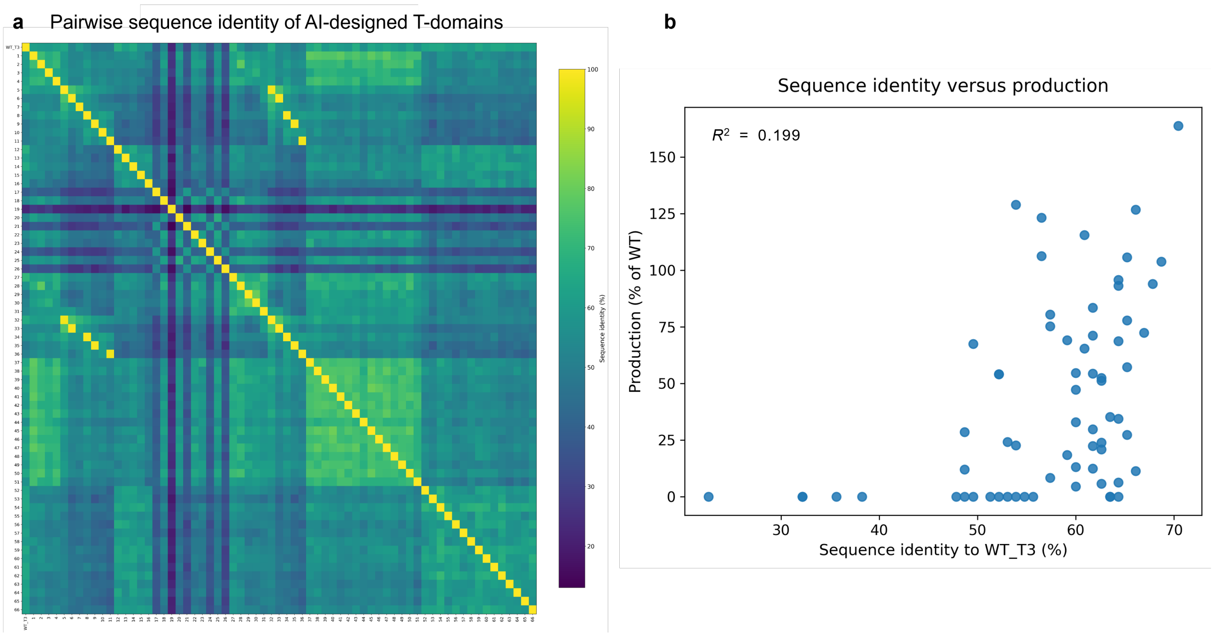

**Supplementary Figure S3.** Sequence similarity landscape and functional performance of AI-designed T-domains. **a,** Pairwise amino acid sequence identity heatmap of the WT GxpS_T3 and 66 AI-designed T-domain variants. Sequence identity values were calculated by global pairwise alignment and are shown as percentages. Variants are ordered consistently along both axes to facilitate comparison across designs. **b,** Relationship between sequence identity to WT GxpS_T3 and peptide production levels for AI-designed T-domain variants in the SU2-only tripeptide assay system ($R^2$ = 0.199). Production is reported as normalized titers relative to the WT GxpS_T3 control. The WT GxpS_T3 sequence itself is excluded from this plot. Each point represents an individual T-domain variant. The lack of a strict correlation highlights that high catalytic performance can be achieved despite substantial sequence divergence from the native T-domain. Source data are provided with this paper.

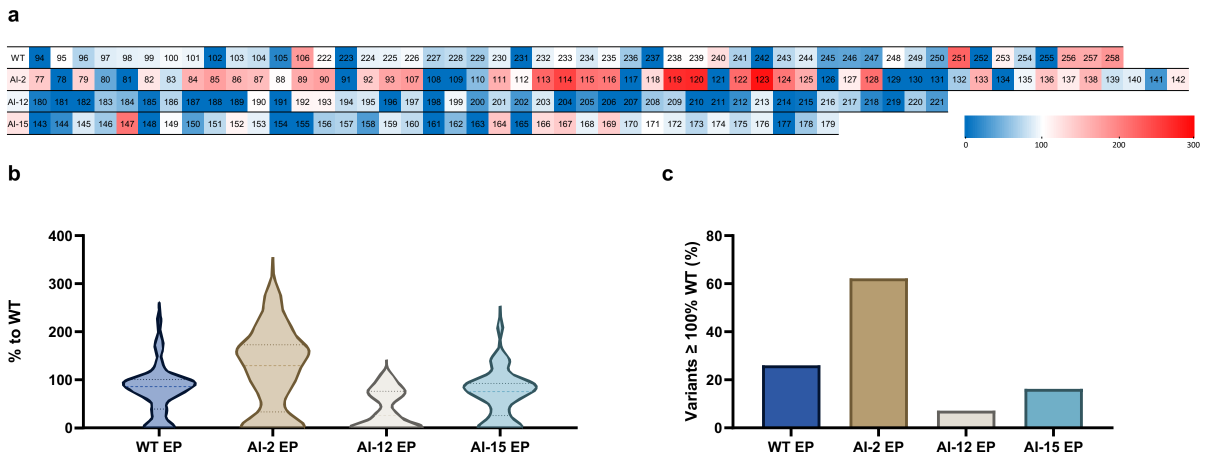

**Supplementary Figure S4.** Performance of error-prone libraries derived from wild-type and AI-designed T-domains. **a**, Heatmap representation of normalized tripeptide production levels for individual variants from the WT GxpS_T3, AI-2, AI-12, and AI-15 error-prone libraries. Each cell represents an individual variant, and the numbers shown correspond to variant IDs. The color scale represents normalized production values ranging from 0 to 300, with blue indicating lower and red indicating higher production levels. **b,** Violin plots of normalized peptide production levels for error-prone variants derived from each parental T-domain group. **c**, Fraction of variants with production ≥ WT, defined as variants exceeding 100 % of WT normalized production. Sample sizes were as follows: WT EP (n = 50), AI-2 EP (n = 53), AI-12 EP (n = 42), AI-15 EP (n = 37). Source data are provided with this paper.

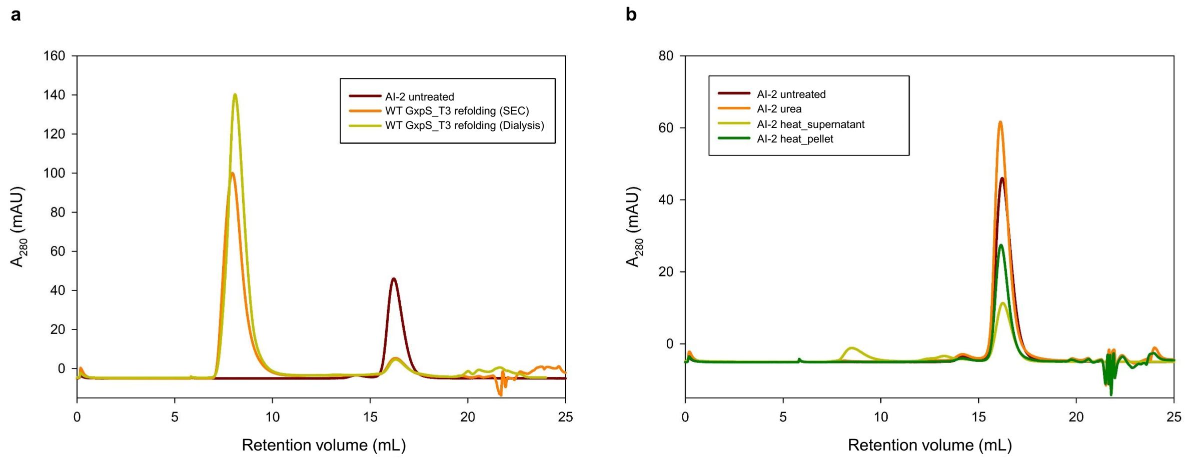

**Supplementary Figure S5.** SEC profiles of WT GxpS_T3 and AI-2. **a,** SEC chromatograms of WT GxpS_T3 following refolding after purification under denaturing conditions, showing a major void-volume peak and a minor species at the elution volume corresponding to the soluble reference. **b,** SEC chromatograms of AI-2 after denaturation in 6 M urea and refolding by SEC, compared to untreated soluble AI-2. Additional traces show SEC profiles of the soluble fraction after heat treatment (95 °C, 10 min; rapid cooling) and of the clarified insoluble fraction. UV absorbance at 280 nm is plotted against retention volume.

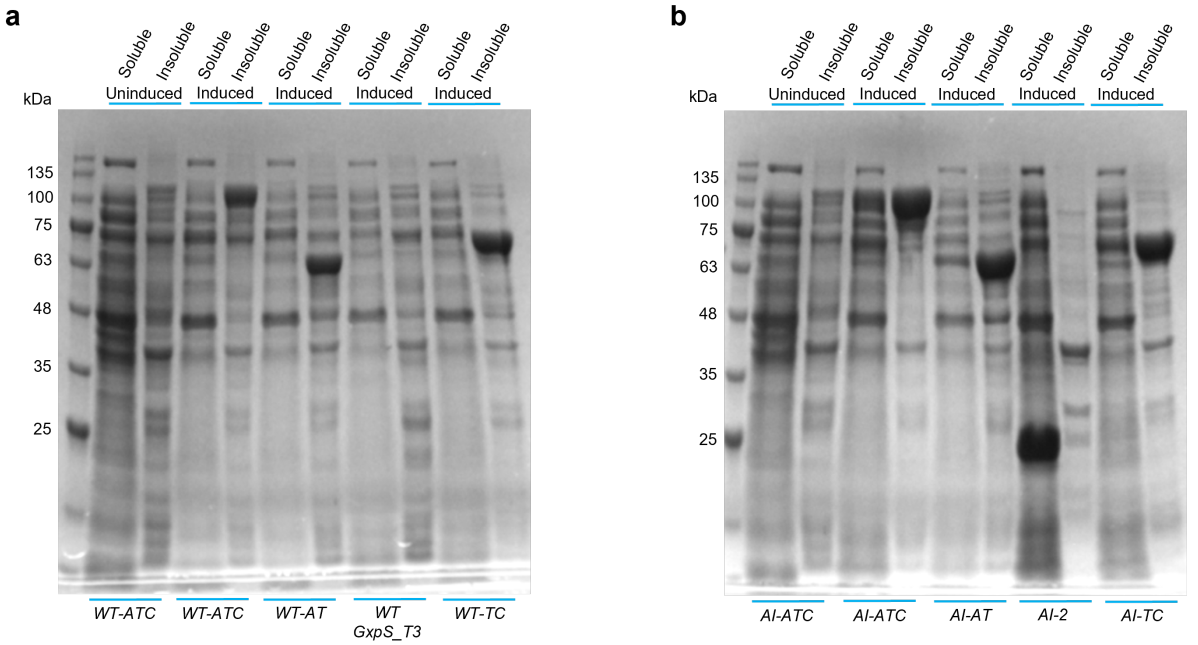

**Supplementary Figure S6**. Expression and solubility of tri-domain and di-domain constructs containing **(a)** WT GxpS_T3 and **(b)** AI-2. Protein expression was induced by addition of 0.2 mM IPTG followed by incubation at 18 °C for 16 hours.

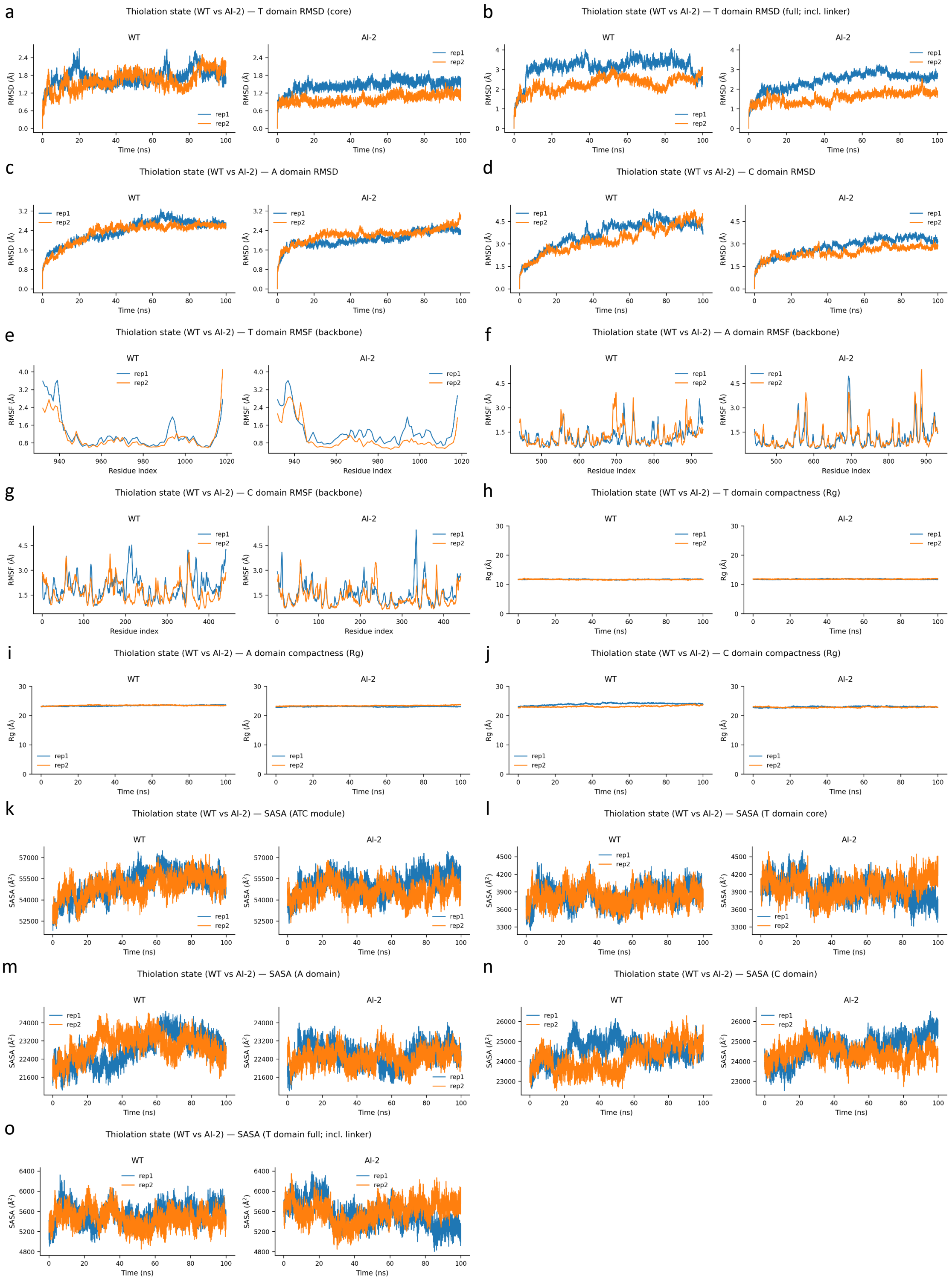

**Supplementary Figure S7.** Structural dynamics of the thiolation state in WT and AI-2 constructs. Molecular dynamics trajectories (100 ns) of the thiolation state were analyzed for the wild-type (WT) and AI-2 variants using two independent replicates per system. Left panels show WT and right panels show AI-2. A: T-domain RMSD without linker. B: T-domain RMSD including linker. C: RMSD of the A-domain. D: RMSD of the C-domain. E: RMSF of the T-domain. F: RMSF of the A-domain. G: RMSF of the C-domain. H: Rg of the T-domain. I: Rg of the A-domain. J: Rg of the C-domain. K: SASA of the ATC tridomain. L: SASA of the T-domain without linker. M: SASA of the A-domain. N: SASA of the C-domain. O: SASA of the T-domain including linker. Time-resolved RMSD are reported for the A-domain, C-domain, T-domain core and full T-domain including the linker, indicating overall structural stability. RMSF of backbone atoms highlight residue-level flexibility across the three domains. Rg profiles for each domain reflect changes in global compactness during the simulation. SASA is shown for the ATC module, individual domains and the T-domain core (and full T-domain where indicated), reporting exposure to solvent over time. For time-dependent analyses, 10,000 frames correspond to 100 ns. WT and AI-2 panels share identical y-axis scaling to enable direct quantitative comparison. No averaging was performed; individual replicate trajectories are shown.

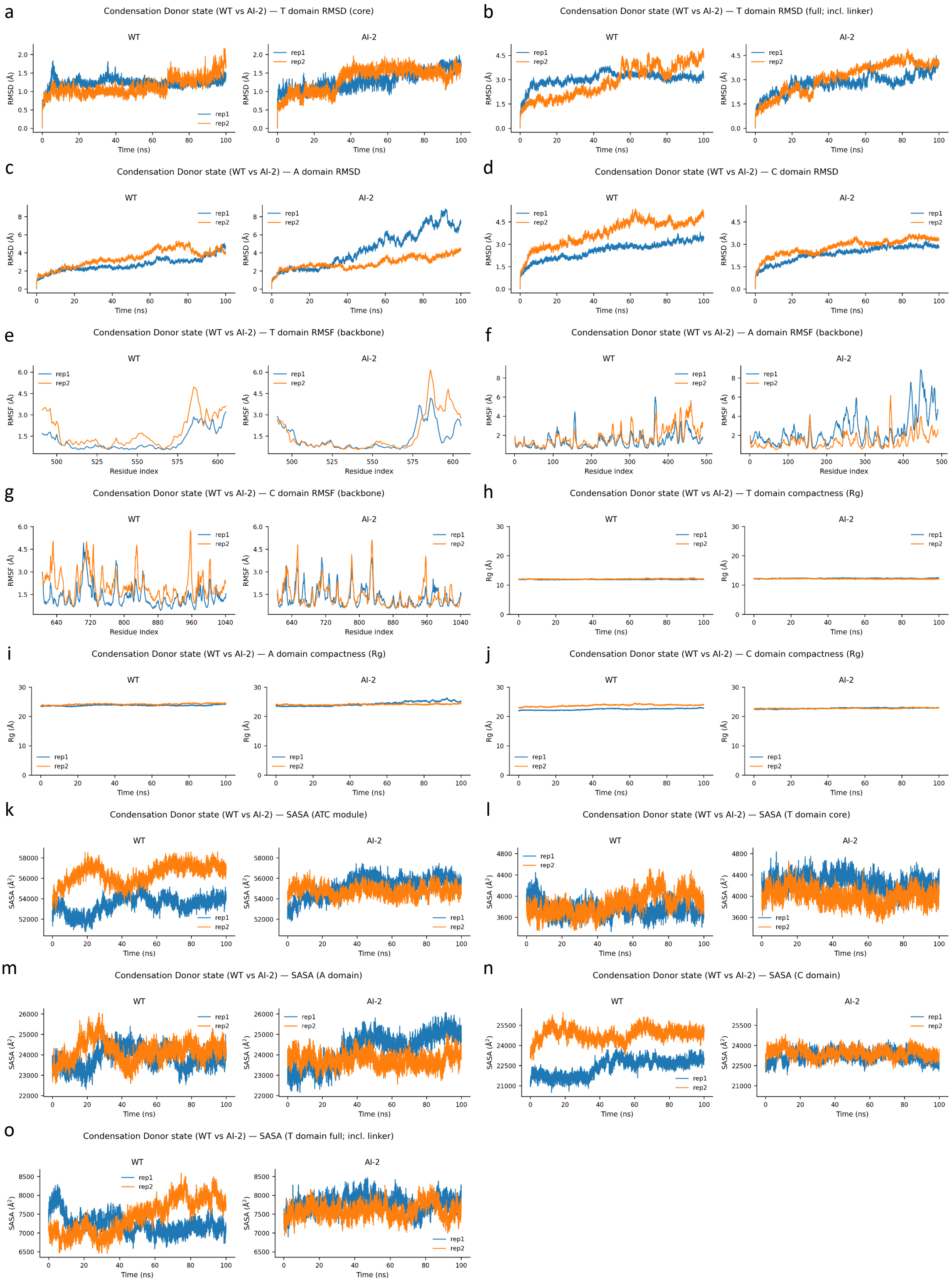

**Supplementary Figure S8.** Structural dynamics of the condensation donor state in WT and AI-2 constructs. Molecular dynamics trajectories (100 ns) of the condensation donor state were analyzed for the wild-type (WT) and AI-2 variants using two independent replicates per system. Left panels show WT and right panels show AI-2. A: T-domain RMSD without linker. B: T-domain RMSD including linker. C: RMSD of the A-domain. D: RMSD of the C-domain. E: RMSF of the T-domain. F: RMSF of the A-domain. G: RMSF of the C-domain. H: Rg of the T-domain. I: Rg of the A-domain. J: Rg of the C-domain. K: SASA of the ATC tridomain. L: SASA of the T-domain without linker. M: SASA of the A-domain. N: SASA of the C-domain. O: SASA of the T-domain including linker. Time-resolved RMSD are reported for the A-domain, C-domain, T-domain core and full T-domain including the linker, indicating overall structural stability. RMSF of backbone atoms highlight residue-level flexibility across the three domains. Rg profiles for each domain reflect changes in global compactness during the simulation. SASA is shown for the ATC module, individual domains and the T-domain core (and full T-domain where indicated), reporting exposure to solvent over time. For time-dependent analyses, 10,000 frames correspond to 100 ns. WT and AI-2 panels share identical y-axis scaling to enable direct quantitative comparison. No averaging was performed; individual replicate trajectories are shown.

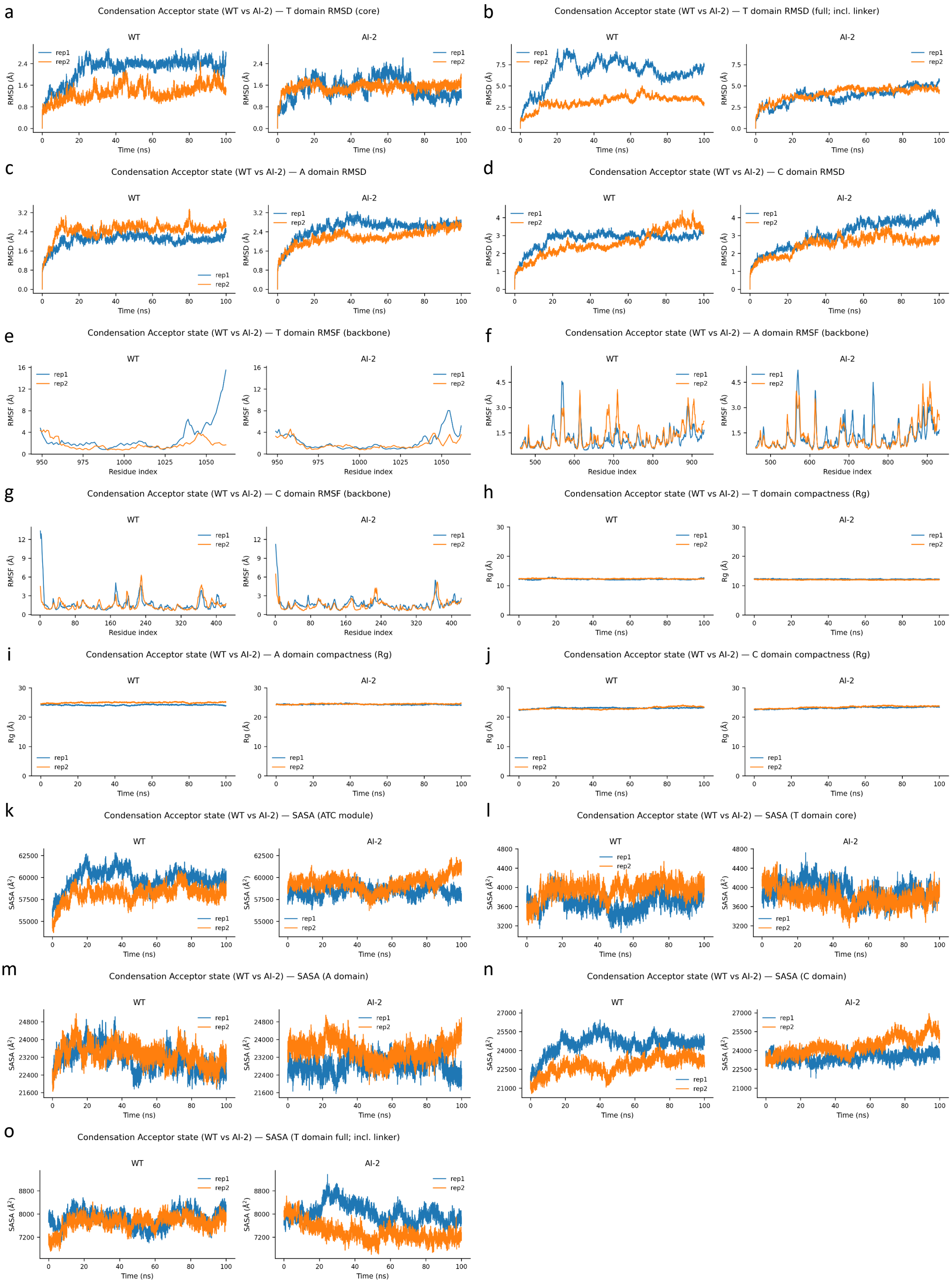

**Supplementary Figure S9.** Structural dynamics of the condensation acceptor state in WT and AI-2 constructs. Molecular dynamics trajectories (100 ns) of the condensation acceptor state were analyzed for the wild-type (WT) and AI-2 variants using two independent replicates per system. Left panels show WT and right panels show AI-2. A: T-domain RMSD without linker. B: T-domain RMSD including linker. C: RMSD of the A-domain. D: RMSD of the C-domain. E: RMSF of the T-domain. F: RMSF of the A-domain. G: RMSF of the C-domain. H: Rg of the T-domain. I: Rg of the A-domain. J: Rg of the C-domain. K: SASA of the ATC tridomain. L: SASA of the T-domain without linker. M: SASA of the A-domain. N: SASA of the C-domain. O: SASA of the T-domain including linker. Time-resolved RMSD are reported for the A-domain, C-domain, T-domain core and full T-domain including the linker, indicating overall structural stability. RMSF of backbone atoms highlight residue-level flexibility across the three domains. Rg profiles for each domain reflect changes in global compactness during the simulation. SASA is shown for the ATC module, individual domains and the T-domain core (and full T-domain where indicated), reporting exposure to solvent over time. For time-dependent analyses, 10,000 frames correspond to 100 ns. WT and AI-2 panels share identical y-axis scaling to enable direct quantitative comparison. No averaging was performed; individual replicate trajectories are shown.

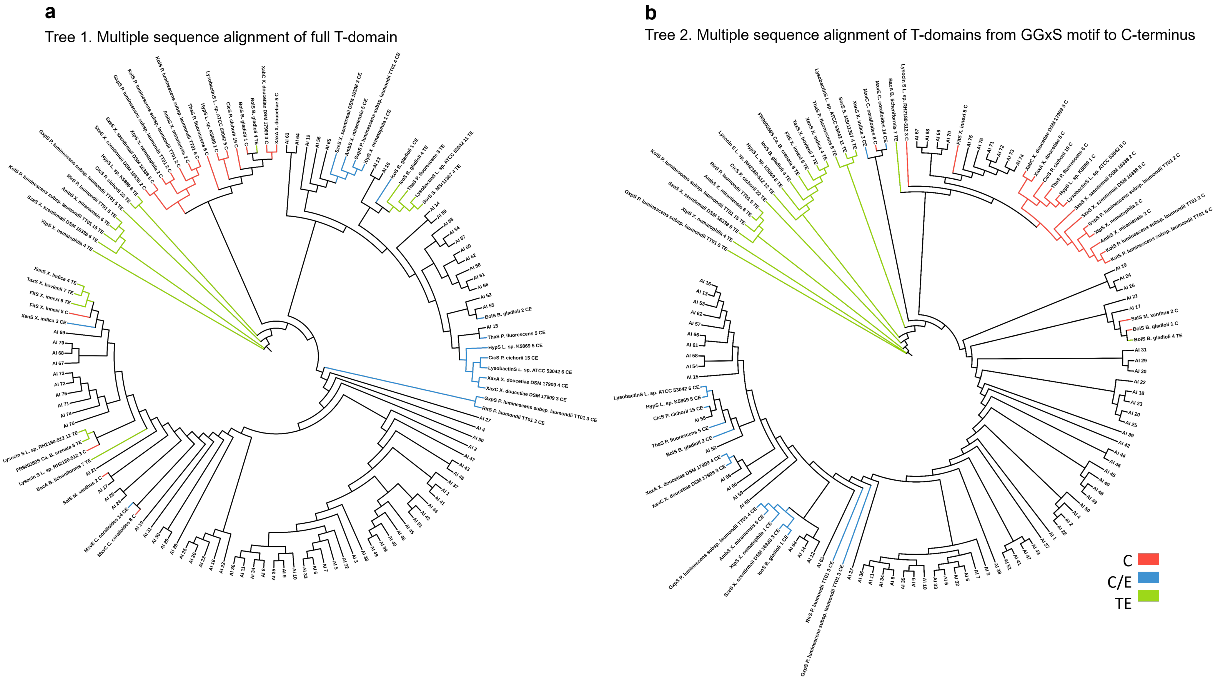

**Supplementary Figure S10.** Phylogenetic analysis of AI-designed and reference T-domains with downstream context. **a**, Tree 1 inferred from the full-length T-domain. **b**, Tree 2 inferred from the alignment positions beginning at the conserved GGXS motif (within FFXXGGXS sequence) to the C terminus, corresponding to the downstream-facing region of the T-domain. AI-generated T-domain sequences are labeled as AI-n, where n denotes the variant identifier assigned during the design workflow, whereas native NRPS sequences are annotated with the NRPS name, taxon name, module number, and the downstream domain. T-domains are color-coded by the downstream domain type: C (red), C/E (blue), and TE (green).

a

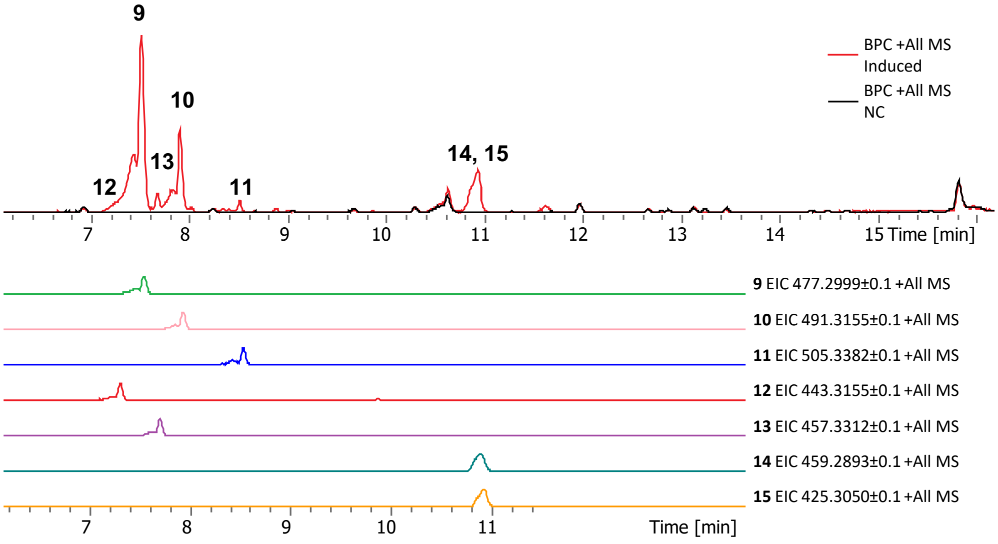

b

vLfV (m/z 477.3071, [M+H]$^+$)

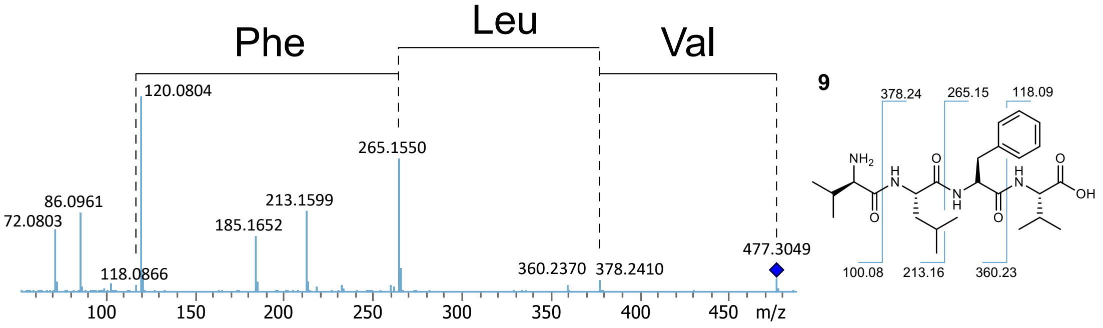

c

lLfV (m/z 491.3228, [M+H]$^+$)

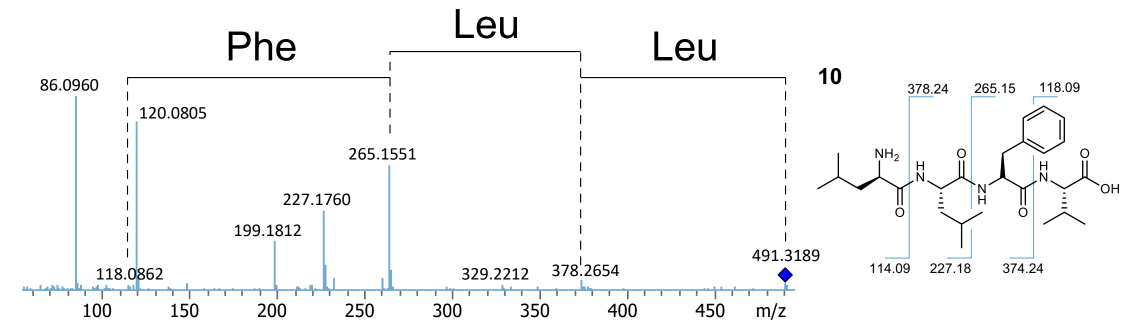

**d**

vLfV-OEt (m/z 505.3384, [M+H]$^+$)

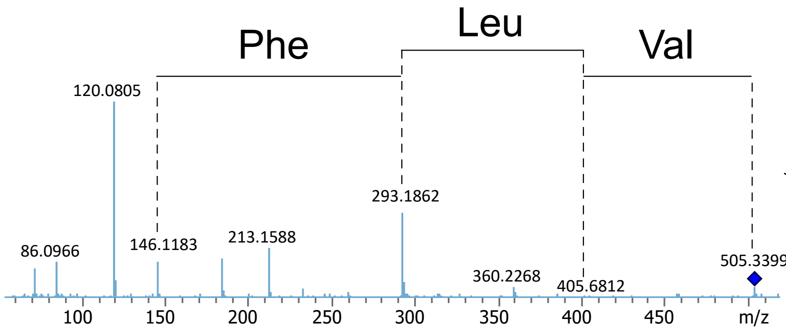

**11**

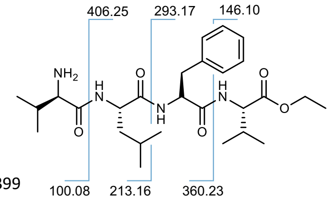

**e**

vLlV (m/z 443.3228, [M+H]$^+$)

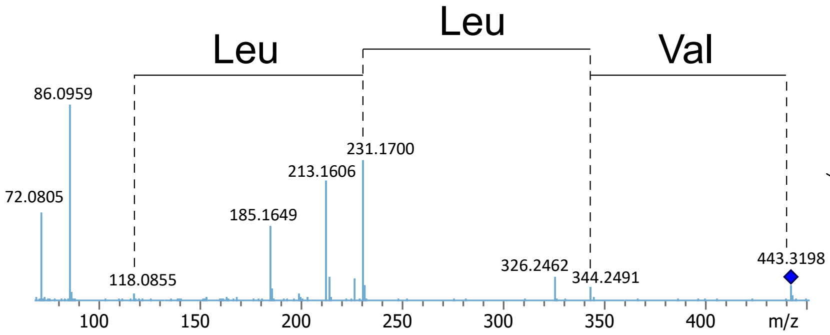

**12**

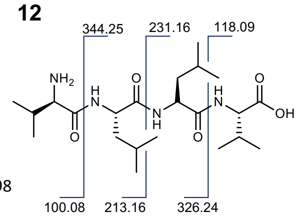

**f**

lLlV (m/z 457.3384, [M+H]$^+$)

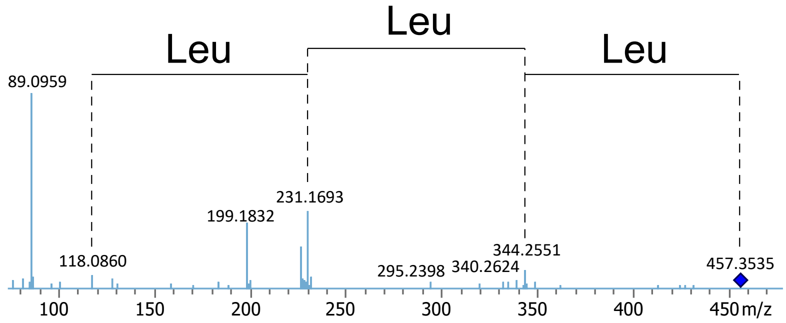

**13**

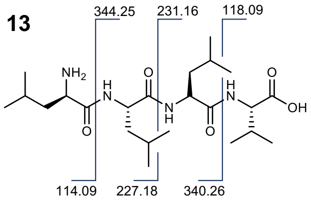

**g**

cyclo vLfV (m/z 459.2966, $[M+H]^+$)

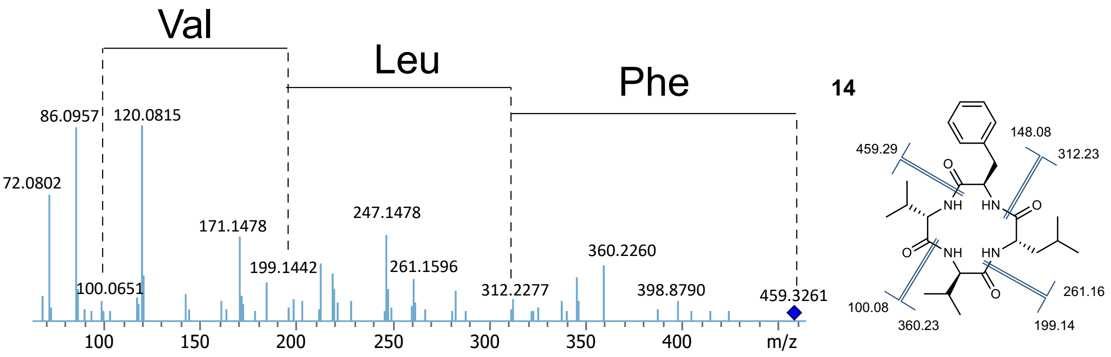

**h**

cyclo vLlV (m/z 425.3122, $[M+H]^+$)

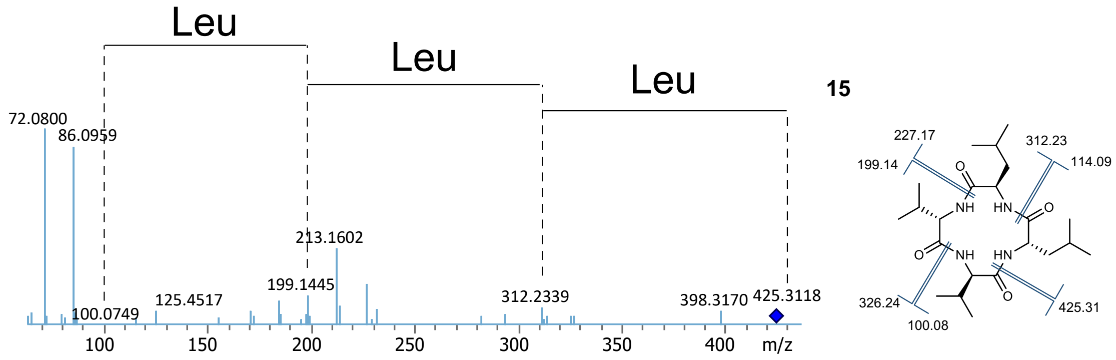

**Supplementary Figure S11. a,** Base peak chromatograms (BPCs) and extracted ion chromatograms (EICs) of GxhS-1 expressed peptides. Distinct peaks corresponding to peptide products are numbered as **9–15**. **b–h**, MS/MS spectrum and structural annotations of **9–15**.

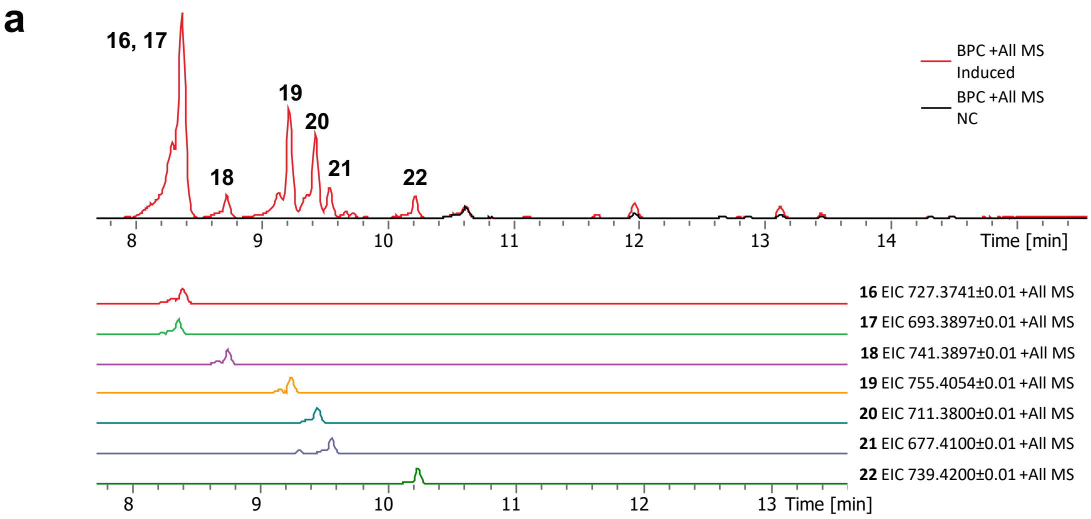
a
16, 17
19
20
18
21
22
BPC +All MS Induced
BPC +All MS NC
Time [min]
16 EIC 727.3741±0.01 +All MS
17 EIC 693.3897±0.01 +All MS
18 EIC 741.3897±0.01 +All MS
19 EIC 755.4054±0.01 +All MS
20 EIC 711.3800±0.01 +All MS
21 EIC 677.4100±0.01 +All MS
22 EIC 739.4200±0.01 +All MS
Time [min]

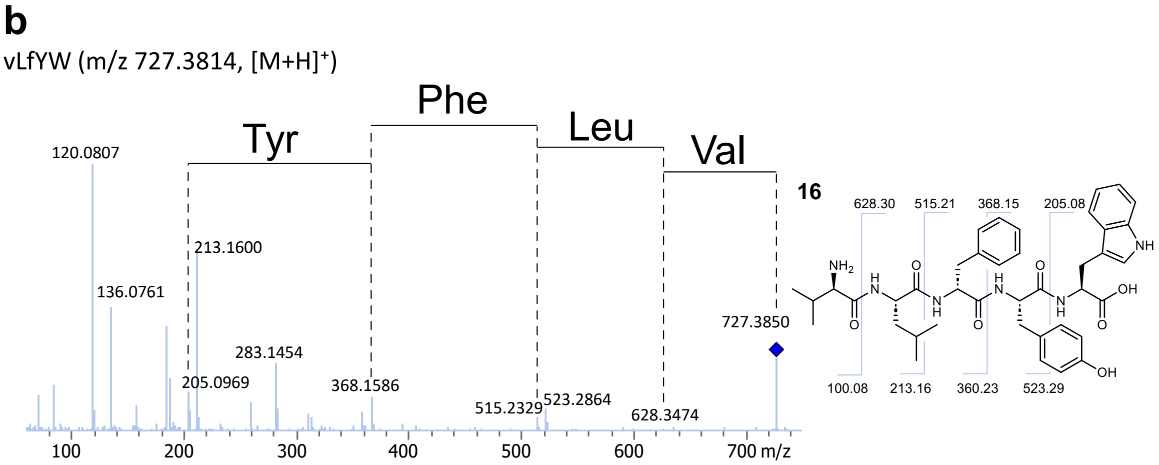
b
vLfYW (m/z 727.3814, [M+H]+)
Tyr
Phe
Leu
Val
120.0807
213.1600
136.0761
283.1454
205.0969
368.1586
515.2329
523.2864
628.3474
727.3850
m/z
16
628.30
515.21
368.15
205.08
100.08
213.16
360.23
523.29

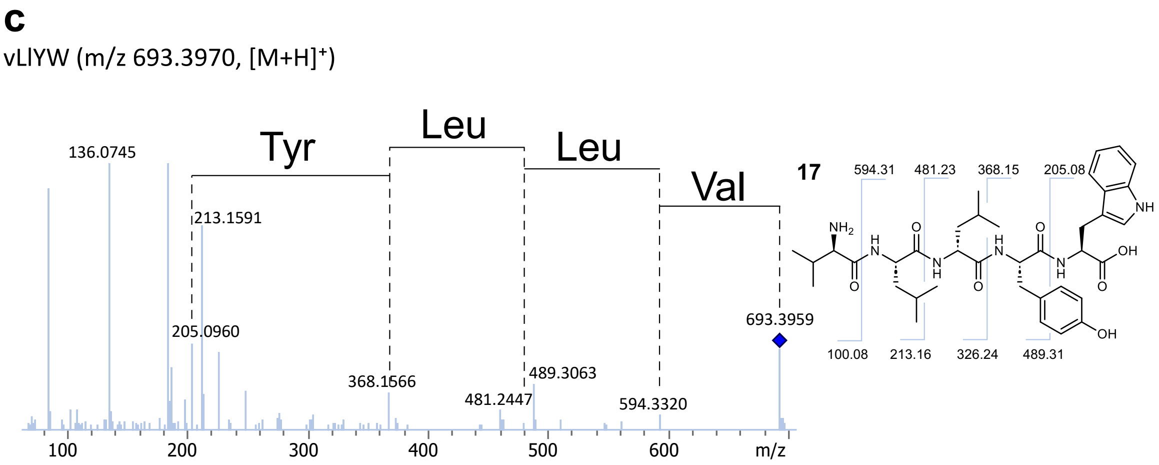
c
vLlYW (m/z 693.3970, [M+H]+)
Tyr
Leu
Leu
Val
136.0745
213.1591
205.0960
368.1666
481.2447
489.3063
594.3320
693.3959
m/z
17
594.31
481.23
368.15
205.08
100.08
213.16
326.24
489.31

**g**

vLlFW (m/z 677.4021, [M+H]⁺)

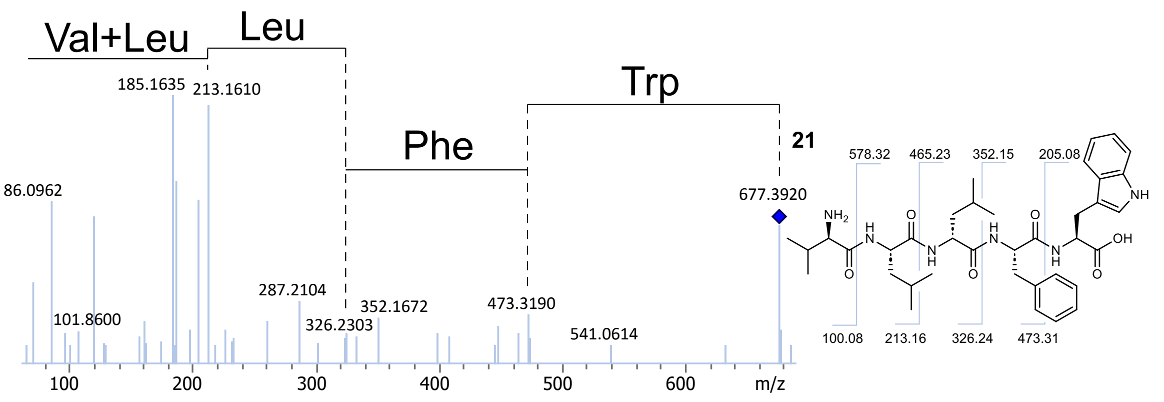

**h**

vLfFW-OEt (m/z 739.4178, [M+H]⁺)

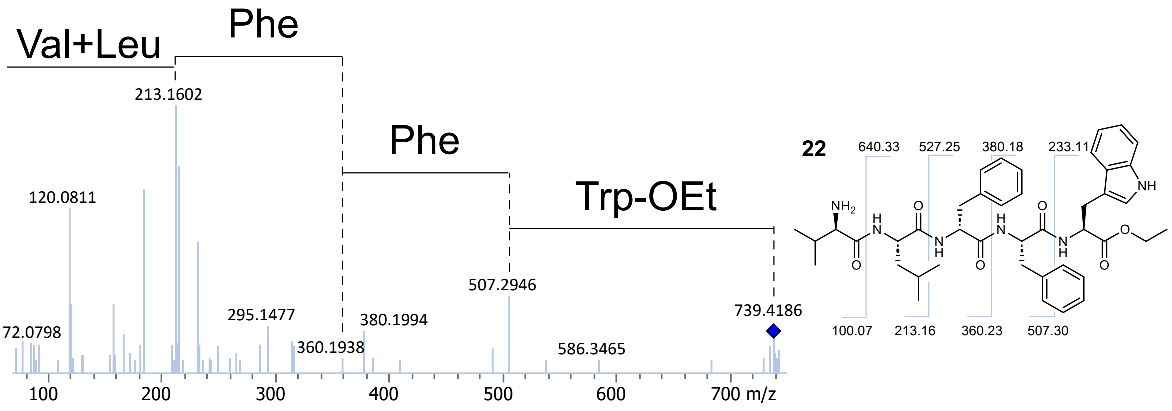

**Supplementary Figure S12**. **a,** Base peak chromatograms (BPCs) and extracted ion chromatograms (EICs) of GshS expressed peptides. Distinct peaks corresponding to peptide products are numbered as **16–22**. **b–h**, MS/MS spectrum and structural annotations of **16– 22**.

**a**

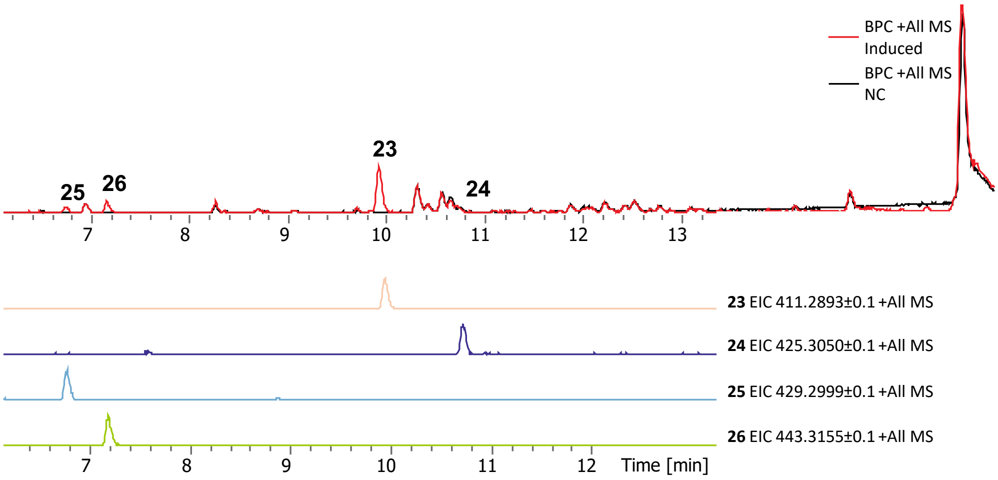

**b**

cyclo vLvV (m/z 411.2966, [M+H]$^+$)

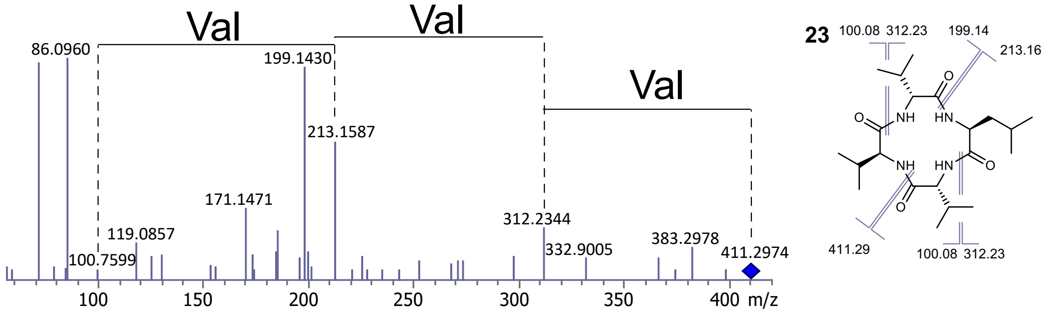

**c**

cyclo lLvV (m/z 425.3122, [M+H]$^+$)

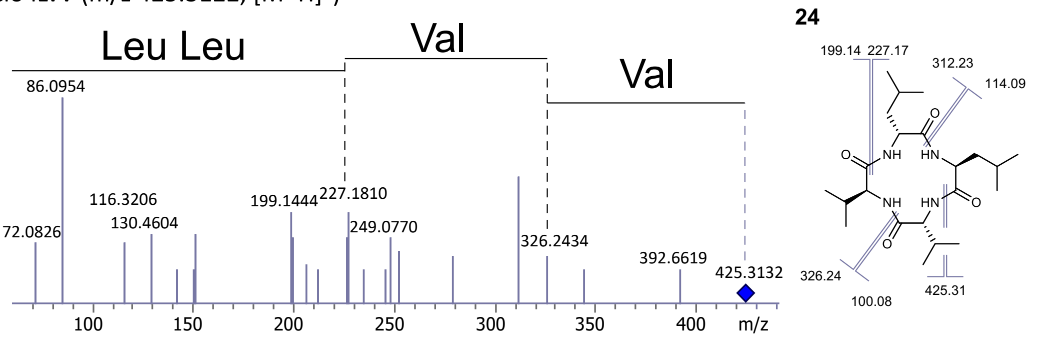

**d**

vLvV (m/z 429.3071, [M+H]$^+$)

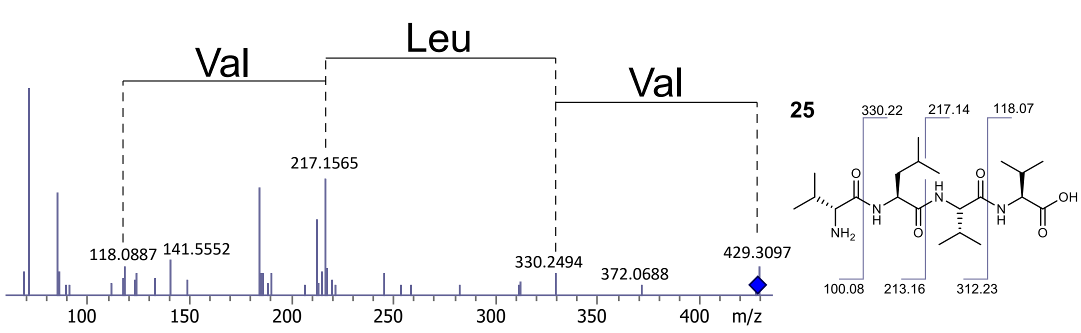

**e**

lLvV (m/z 443.3228, [M+H]⁺)

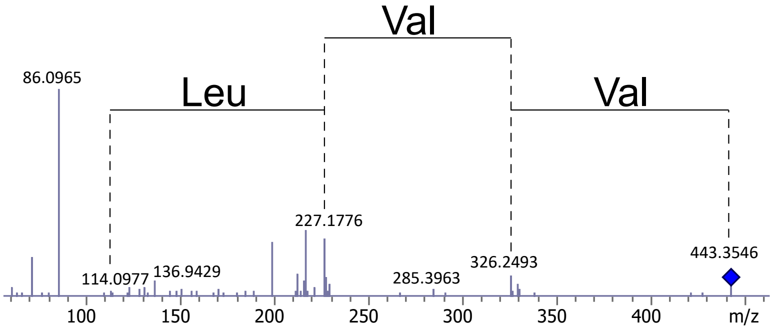

**26**

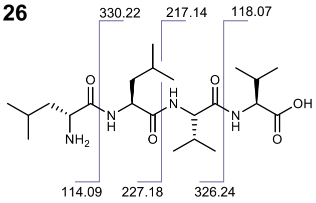

**Supplementary Figure S13**. **a,** Base peak chromatograms (BPCs) and extracted ion chromatograms (EICs) of GxhS-2 expressed peptides. Distinct peaks corresponding to peptide products are numbered as **23–26. b - e**, MS/MS spectrum and structural annotations of **23–26**.

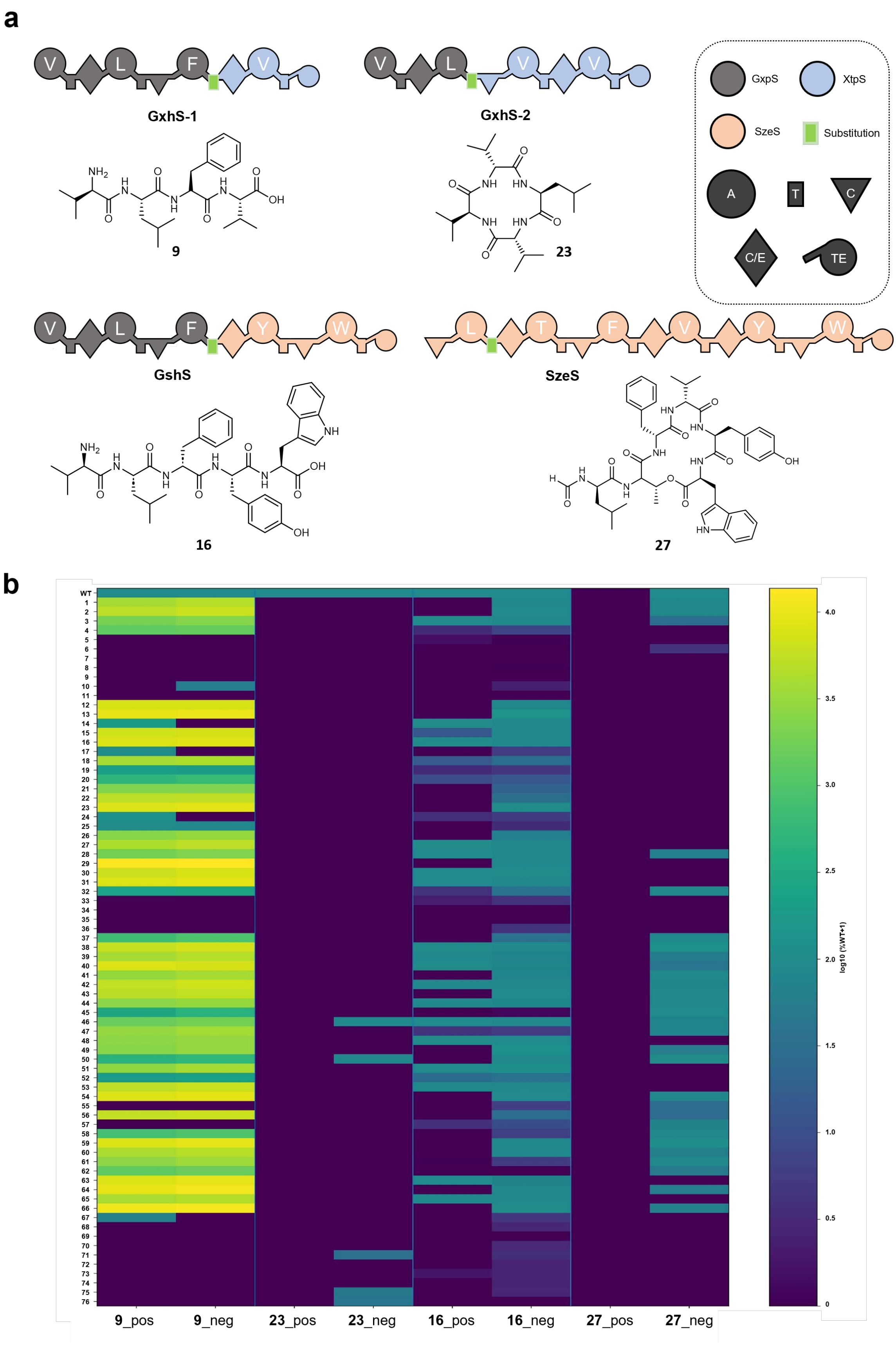

**Supplementary Figure S14**. RapidFire LC–MS screening of T-domain variants across GxhS-1, GxhS-2, GxhS, and SzeS systems. **a,** Schematic representation of the native and hybrid NRPS architecture used for screening (GxhS-1, GxhS-2, GxhS, and SzeS) and the corresponding major peptide products detected in each system. **b**, Heatmap shows WT-normalized product formation (%WT) for the main product detected in each system: vLfV (**9**) (GxhS-1), cyclo(vLvV) (**23**) (GxhS-2), vLfYW (**16**) (GxhS), and cyclo(lTfvYW) (**27**) (SzeS), across all T-domain variants. Only the dominant product in each system is shown; additional minor side products detected by RapidFire LC–MS are summarized in Supplementary Data 1. Positive- and negative-ionization mode peak heights were normalized independently to the corresponding wild-type (WT) signal for each product (WT = 100 %) after background subtraction and setting negative values to zero. For SzeS, the WT signal was detected only in negative ionization mode; therefore, normalization and visualization are shown exclusively for the negative mode, and positive-mode data are omitted. Columns represent system–ionization-mode pairs (e.g., GxhS vLfYW_positive_mode and GxhS vLfYW_negative_mode), and rows represent T-domain variants labeled by variant number (y-axis). The color scale represents log10(%WT + 1) to compress dynamic range while preserving quantitative relationships (100 % WT corresponds to log10(101) ≈ 2.0). Values of 0 indicate no detectable product above the background. Source data are provided with this paper.

#### Supplementary Note 1: AI workflow for generation of T-domains

##### Library generation using generative protein models

T-domain sequences were generated using three pretrained generative models: ESM3[5], EvoDiff[6] and ProteinMPNN[7]. For all models, we followed a conditional sequence design strategy where the amino acid identities of the flanking adenylation (A) and condensation (C) domains were kept fixed and used as context to generate the masked T-domain in the middle. This ensured that each method produced T-domain variants explicitly conditioned on the surrounding domain context.

##### ProteinMPNN

We used the openly available 0.02 Å ProteinMPNN model (*v_48_002,* https://github.com/dauparas/ProteinMPNN/tree/main/vanilla_model_weights) for T-domain design, chosen for its minimal structural perturbation. Empirically, this version performed best among the ProteinMPNN variants for our application. We set the sampling temperature to 0.3 to balance sequence diversity and sequence recovery. The remaining parameters were kept at their default values. We provided the A-T-C tri-domain structure, created using homology modelling via SWISS-MODEL[8] based on crystal structures (PDB IDs 6MFY[9], 6MFZ[9], and 6MG0[9]), as input to the model. During decoding, the amino acid residues of the A- and C-domains were fixed, ensuring that only the T-domain was generated.

##### ESM3

ESM3 is a multi-modal large language model that can reason across sequence, structure, and function modalities. We used the publicly available (https://github.com/evolutionaryscale/esm) 1.4B parameter version for our design tasks. Two types of designs were generated. In the first setting, both sequence and structure tracks were used for T-domain generation. For this, we provided the A-T-C tri-domain sequence with the T-domain region masked using the mask token together with its 3D structure. These designs are labelled with _str_ (see Supplementary Data). In the second setting, only the sequence track was used. A decoding temperature of 0.5 was used for all generations. Decoding was performed iteratively, and the number of steps was set to half the length of the T-domain.

##### EvoDiff

We used the publicly available (https://github.com/microsoft/evodiff) 640M parameter order-agnostic autoregressive diffusion (OADM) model. The A-T-C tri-domain sequence was provided as input, with the T-domain region masked. All parameters were kept at their default values.

##### *In silico* evaluation

To evaluate the generated T-domains, we assess the sequences using multiple *in silico* metrics. Detailed descriptions are provided below.

**Sequence Identity:** This metric provides a measure of similarity between the generated sequences and the wildtype T-domain. To calculate sequence identity, we first align the two sequences using *hmmalign* from HMMER[10]. The HMM profile for the T-domain was retrieved from InterPro (PFAM Entry PF00550)[11]. Using sequence identity as an evaluation criterion, we assess the ability of the models to generate plausible T-domain sequences for the given context. Higher sequence identity values suggest that a model produces sequences that are more structurally and functionally aligned with the wildtype.

**Foldability and Structural Similarity:** To assess structural plausibility, we refold the generated sequences and evaluate their predicted structures. We use ESMFold[12] to predict the structures of the designed T-domains.

- For foldability, we use the pLDDT scores from ESMFold. pLDDT reflects the confidence of the model in the predicted structure at the residue level. For each generated sequence, we calculate the mean pLDDT as a measure of foldability.
- For structural similarity, we compare the predicted structure to the ground-truth structure. We use TM-align to compute the TM-score[13,14]. Higher TM-scores indicate that the generated sequences fold into structures closely resembling the original, potentially retaining similar functionality.

**Perplexity:** Perplexity is commonly used in natural language processing (NLP) to evaluate how well a probabilistic model predicts a sequence[15]. In our design setting, we use perplexity to assess the biological plausibility of the T-domain sequences generated by our models. We use the 650M ESM2 model[12] to calculate perplexity. We estimate perplexity as follows:

$$perplexity = exp\left(-\frac{1}{L}\sum_{i=1}^{L} log\, p(x_i|x_{-i})\right)$$

where $L$ is the length of the amino acid sequence and $p(x_i|x_{-i})$ is retrieved from the softmax probabilities of ESM2 at position $i$, when position $i$ is masked.

**Surrogate Models**

For round 3, we implemented a surrogate model to guide the selection of candidate T-domains for experimental testing. We used the T-domain sequences and their corresponding activity values from rounds 1 and 2 as the training set. In total, this dataset comprised 85 labeled sequences evaluated in the SU2-only GxpS assay, using fIL (**1**) and FIL (**2**) production as the quantitative fitness readout. We evaluated multiple model architectures based on their ability to rank T-domain sequences. Specifically, we considered three categories: (i) models using pretrained protein language model (PLM) embeddings as protein representations, (ii) zero-shot fitness scores derived from PLMs, and (iii) fine-tuned PLMs. We used the ESMC 600M[16] as the base PLM. We also evaluated ESM2, but ESMC consistently provided better performance.

For models using pretrained representation, we adopted two strategies to obtain the final representation: (i) mean pooling of embeddings across the residue dimension, and (ii) unrolling the residue-level embeddings into a single concatenated vector. On top of these representations, we evaluated three prediction heads: a linear regression head, a shallow multi-layer perceptron (MLP), and a random forest regressor. In all cases, only the prediction head was trained, while the PLM parameters remained frozen. Models were trained to predict T-domain activity using mean squared error (MSE) loss.

For mean-pooled embeddings, the random forest and shallow MLP achieved comparable performance and outperformed the linear head. We therefore selected the random forest for subsequent experiments. This motivated both by its strong empirical performance and by its successful application in the recent EvolvePro[17] framework. In contrast, for the unrolled embedding representation, a linear head performed best and was used in subsequent experiments. We hypothesize that the random forest and MLP were more prone to overfitting in this high-dimensional setting, whereas the simpler linear model generalized better.

Meier et al. (2021) demonstrated that PLM-derived likelihoods can be used for zero-shot fitness prediction and outlined multiple approaches for scoring protein variants based on these likelihoods[18]. In this study, we used the masked marginal score, estimated by computing the log-odds ratio of the mutant amino acid relative to the wildtype amino acid at each mutated position. Let $M$ denote the set of mutated positions in a variant relative to the wildtype sequence. The masked marginal fitness score is defined as:

$$\widehat{f_\theta}(x) = \sum_{i \in M} log\, p(x_i^{mt} | x_{-M}) - log\, p(x_i^{wt} | x_{-M})$$

where $x_{-M}$ denotes the sequence with masked tokens at the mutated positions, and $x_i^{mt}$ and $x_i^{mt}$ represent the amino acid identities at position $i$ in the mutant and wildtype sequences, respectively.

For fine-tuning ESMC, we adopted two strategies. The first followed a conventional setup, in which we extracted the class token (<CLS>) embedding and applied a linear layer to predict activity values[19]. To enable efficient fine-tuning and reduce overfitting, we use Low-Rank Adaptation (LoRA) to update only a fraction of the total model parameters[20]. This model was trained using MSE loss. The second approach employed a contrastive fine-tuning strategy using a ranking objective[21]. The loss function was based on the Bradley–Terry (BT) model, which enforces pairwise ranking consistency. Given two sequences, $x^{(i)}$ and $x^{(j)}$, with corresponding fitness values $y^{(i)}$ and $y^{(j)}$, where $y^{(i)} > y^{(j)}$, the BT loss is defined as:

$$L_{BT} = \sum_{y^{(i)} > y^{(j)}} log \left[ 1 + e^{-\left( \widehat{f_\theta}(x^{(i)}) - \widehat{f_\theta}(x^{(j)}) \right)} \right]$$

where $\widehat{f_\theta}$ represents the fitness function parameterized by $\theta$. We used the masked marginal score described above as the fitness function. To prevent overfitting, we introduced a Kullback–Leibler (KL) divergence regularization term:

$$L_{KL} = D_{KL}(P_\theta || P_{\theta_0}) = \sum_i P_\theta(x_i | x_{-i})\, log \frac{P_\theta(x_i | x_{-i})}{P_{\theta_0}(x_i | x_{-i})}$$

where $P_\theta$ represents the probability distribution of the fine-tuned PLM and $P_{\theta_0}$ corresponds to the distribution of the pretrained PLM. The final training objective is formulated as:

$$L = L_{BT} + \lambda L_{KL}$$

where $\lambda$ is a hyperparameter controlling the contribution of the KL divergence regularization term. LoRA was also used in this setting to fine-tune ESMC.

**Training details**

For models using pretrained embeddings, we first extracted sequence embeddings from the final layer of ESMC for all sequences and stored them. These embeddings were then used as input to downstream predictive models. We implemented the linear model using the Ridge module from scikit-learn (version 1.4.2) with default parameters and random forest using RandomForestRegressor module.

For fine-tuning ESMC in both settings described above, we employed LoRA with rank 8, alpha 8, and a dropout rate of 0.1. LoRA adapters were applied to the key, query, and value projection layers, as well as to the feed-forward network following the Multi-Head Attention block. For contrastive fine-tuning, LoRA was additionally applied to the sequence head and the KL divergence regularization weight (λ) was set to 0.1. Training was performed with early stopping using a patience of 10 epochs. A batch size of 8 was used with gradient accumulation over 4 steps, resulting in an effective batch size of 32. Experiments were conducted using PyTorch version 2.3.1 and PyTorch Lightning version 2.5.

#### Results

The surrogate models were benchmarked using 85 T-domain sequences from rounds 1 and 2. These were split into training (n = 55) and test (n = 30) sets. To minimize data leakage and ensure a rigorous evaluation, we performed the split based on Hamming distance, such that sequences in the test set were maximally distant from those in the training set. In the final split, the nearest test sequence was at least 25 mutations away from any training sequence. Models were trained on the training set, and performance was evaluated on the test set using Spearman rank correlation between predicted and experimentally measured activity (Supplementary Table S4). We see that the unrolled pretrained ESMC embedding with a linear head achieved the highest mean performance, followed by contrastive fine-tuning.

Supplementary Table. S4: Mean values and 95% confidence intervals (CI) for spearman correlation on the T-domain test set. CI was estimated using bootstrapping. RF: Random Forest.

| Model | T-domain |
|---|---|
| Mean pooling + RF head | 0.25 (-0.10, 0.59) |
| Unrolled + linear head | **0.57** (0.26, 0.83) |
| Masked marginal score | 0.35 (-0.05, 0.74) |
| MSE-based finetuning | 0.38 (0.00, 0.72) |
| Contrastive finetuning | 0.43 (0.11, 0.73) |

To have a broader comparison, we further evaluated our models on two widely used protein fitness prediction datasets. First, we used the GB1 dataset, comprising variants of the first binding domain of protein G with one to four mutations at defined positions[22]. We adopted the three-vs-rest split, in which variants containing up to three mutations were used for training and four-mutation variants were reserved for testing[19,23]. Second, we evaluated the TEM1 β-lactamase dataset, which captures the fitness effects of single amino acid substitutions across the protein[24]. Here, we applied a contiguous split scheme[25], dividing the protein sequence into ten segments, with two segments designated as test data and the remaining eight used for training. Results are reported in Supplementary Table S5. On GB1, MSE-based fine-tuning achieved the best performance, whereas contrastive fine-tuning performed best on TEM1. Notably, the masked marginal score performed comparatively well on TEM1 but poorly on GB1. This shows that zero-shot likelihood-based scores may be dataset dependent and that task-specific adaptation, either via fine-tuning or training a predictive head, is necessary for optimal performance.

Supplementary Table. S5: Mean values and 95% confidence intervals (CI) for spearman correlation on both the GB1 and TEM1 dataset. CI was estimated using bootstrapping. RF: Random Forest

| Model | GB1 | TEM1 |
|---|---|---|
| Mean pooling + RF head | 0.5 (0.48, 0.52) | 0.74 (0.71, 0.76) |
| Unrolled + linear head | 0.74 (0.73, 0.76) | 0.84 (0.82, 0.85) |
| Masked marginal score | 0.29 (0.26, 0.31) | 0.82 (0.81, 0.84) |

| | | |
|---|---|---|
| MSE-based finetuning | **0.81** (0.80, 0.82) | 0.71 (0.69, 0.74) |
| Contrastive finetuning | 0.77 (0.76, 0.78) | **0.88** (0.87, 0.89) |

To better mimic the low-data regime of the T-domain setting, we simulated low-data training scenarios on both GB1 and TEM1 while retaining their full test sets for robust evaluation. Specifically, we constructed training subsets of size 50, 100, and 200, sampled such that their fitness distributions closely matched that of the T-domain training set. This analysis enabled us to assess model robustness under limited data and examine scaling behavior with increasing training size (see Supplementary Table S6). In these low-data regimes, MSE-based fine-tuning performed poorly in most settings. In contrast, except for GB1 at N = 200, contrastive fine-tuning consistently outperformed alternative approaches across datasets.

Supplementary Table. S6: Performance on both the GB1 and TEM1 dataset under low-data setting. Each model was trained five times with different random initializations and the reported values correspond to the mean and standard deviation across these five runs. RF: Random Forest.

| Dataset | Models | Low-data Setting | | |
|---|---|---|---|---|
| | | N = 50 | N = 100 | N = 200 |
| GB1 | Mean pooling + RF head | 0.49±0.04 | 0.51±0.05 | 0.56±0.02 |
| | Unrolled + linear head | 0.61±0.05 | 0.7±0.03 | **0.75±0.01** |
| | Masked marginal score | 0.29±0.00 | 0.29±0.00 | 0.29±0.00 |
| | MSE-based finetuning | 0.12±0.03 | 0.12±0.04 | 0.15±0.02 |
| | Contrastive finetuning | **0.63±0.04** | **0.71±0.03** | 0.74±0.03 |
| TEM1 | Mean pooling + RF head | 0.58±0.04 | 0.6±0.01 | 0.68±0.02 |
| | Unrolled + linear head | 0.64±0.01 | 0.68±0.01 | 0.73±0.03 |
| | Masked marginal score | **0.82±0.00** | **0.82±0.00** | **0.82±0.00** |
| | MSE-based finetuning | -0.13±0.05 | -0.09±0.05 | -0.12±0.02 |
| | Contrastive finetuning | 0.78±0.01 | 0.79±0.01 | 0.81±0.01 |

Based on these combined results, we selected contrastive fine-tuning as the primary surrogate model for subsequent rounds. Given the strong performance of the unrolled pretrained embeddings with a linear head, we also retained this approach to provide complementary predictions and increase the robustness of T-domain selection.

**Generation and filtering of Round 3 candidates**

For round 3, we used ESMC with unrolled embeddings and ESMC with contrastive fine-tuning as our surrogate models. For each model, we trained three ensemble members by bootstrapping the training data. To generate a candidate library for round 3, we used ESM3 and EvoDiff to produce 5,000 T-domain sequences conditioned on the given A- and C-domain context.

First, the generated sequences were first filtered using the same *in silico* metrics applied in round 2. We then removed sequences exhibiting high variance across ensemble predictions. In addition, sequences with a masked marginal score lower than −10 were excluded. Although the masked marginal score did not strongly correlate with activity in rounds 1 and 2, we observed that it was effective in filtering out low-performing sequences.

The remaining sequences were divided into five bins based on the fitness scores predicted by each surrogate model. We then identified sequences that were assigned to the same bin by both models. This voting-like mechanism allowed us to prioritize sequences that were consistently placed in similar fitness categories by both surrogates, thereby increasing confidence in their predicted performance. From each bin, we selected sequences with the highest contrastive fine-tuning scores for experimental evaluation. The final set of selected sequences is presented in Supplementary Data 1.

#### Supplementary Methods

##### Peptide synthesis and purification

##### General Remarks

Fmoc-L-Leu-OH, Fmoc-D-Leu-OH, Fmoc-D-Phe-OH, Fmoc-L-Leu-OH, Fmoc-D-Val-OH and O-(7-azabenzotriazol-1-yl)-N,N,N',N'-tetramethyluronium hexafluorophosphate (HATU) were purchased from BLD Pharma. 2-chlorotrityl chloride resin (200-400 mesh, 1% DVB, 0.4-3 mmol/g capacity. *O*-(1*H*-6-Chlorobenzotriazole-1-yl)-1,1,3,3-tetramethyluronium hexafluorophosphate (HCTU), hexafluoroisopropanol (HFIP) and diisopropylethylamine (DIEA) were purchased from abcr GmbH. dichloromethane (DCM) p.a. and dimethylformamide (DMF) p.a. from VWR International, and anhydrous solvents (DMF, DCM) from Sigma Aldrich. NMR analysis 1 H, COSY, HSQC and HMBC spectra were recorded on a Bruker Avance Neo 500 MHz (1H, 500 MHz; 13C, 126 MHz) with prodigy cryoprobe system. Chemical shifts were recorded as δ values in ppm units and referenced against the residual solvent peak (DMSO-d6, δ= 2.50, 39.52 and CDCl3-d1: δ=7.26, 77.16). High-resolution mass spectra were recorded on a ThermoFisher Scientific (TF, Dreieich, Germany) Q Exactive Focus system equipped with heated electrospray ionization (HESI)-II source. Syntheses were carried out manually in PP syringes equipped with PE frits and PTFE stopcocks (Roth Selection).

##### Peptide synthesis

##### General procedure 1: Determination of amino acids loading on the resin

Fmoc-L-Leu-OH was loaded on 2CT resin was determined by spectrophotometric quantification at λ=301 nm of the dibenzofulvene liberated after deprotection of the Fmoc group with piperidine. To a precisely weighed amount of resin (ca. 5 mg) 1 mL of a solution of piperidine/DMF (2:8) was added. After 20 min under shaking, 100μL were transferred in a quartz cuvette and the volume made up to 1 mL with DMF. The UV absorbance was measured at λ=280 nm against a control of a solution of 100 μL piperidine/DMF (2:8) diluted with DMF to a volume of 1 mL. loading was calculated by $L= 1000*D*(A_s-A_b)*V/(\varepsilon^{290}*m)*l$, where D is the dilution factor, $A_s$ the absorbance of the sample at 290 nm and $A_b$ the absorbance of the blank at 280 nm, V is the volume, $\varepsilon^{290}$ is a molar extinction coefficient of dibenzofulvene at 290 nm ($L\ mol^{-1}\ cm^{-1}$) and l is the path length of the cuvette.

##### General procedure 2: Fmoc deprotection

The resin was agitated for 20 min with a solution of piperidine/DMF (2:8), after which the supernatant was removed. The resin was washed subsequently with DMF (3x), methanol (MeOH) (1x) and DCM (3x).

##### General Procedure 3: Coupling of amino acids to resin-bound peptide

To a solution of Fmoc Protected amino acids (3 eq.) and DIEA (6 eq.) in DMF (0.2M), HCTU (4 eq.) was added. The mixture was agitated for 2 min and subsequently added to the pre-swollen resin in DMF. The resin was agitated at rt for 1h, after which the supernatant was removed (completion of coupling checked by treating a small sample of the resin with HFIP to obtain a sample peptide for HPLC/MS analysis) and the resin washed DMF (3x), MeOH (1x) and DCM (3x).

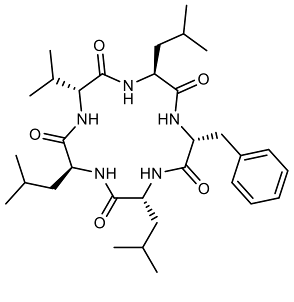

**(3R,6S,9R,12S,15R)-3-benzyl-6,12,15-triisobutyl-9-isopropyl-1,4,7,10,13-pentaazacyclopentadecane-2,5,8,11,14-pentaone (6)**

HRMS (ESI) m/z: calculated for $[M+H]^+$ : 586.3963; found: 586.3925.

For the synthesis of **6**, Fmoc–SPPS procedure was adapted from the literature.[1] To 1 gr of 2CT resin, 10 mL of DCM were added and agitated for 30 min. The solvent was drained off, and a suspension of Fmoc-L-Leu-OH (1.6 gr, 3 equiv.) and DIEA (1.5 mL, 6 equiv.) in DCM (10 mL) was added to the resin. The resin was shaken overnight, drained off and washed 3xDCM and dried. The loading was calculated as General Procedure 1 (0.75 mmol/gr). The sequent deprotection and coupling steps were carried out following General procedure 2 and General Procedure 3.

After completing the amino acids sequence, the resin was suspended in 1:1 DCM/HFIP (20 mL) solution and agitated for 1h. This step was repeated twice. The solution was drained off in a flask and evaporated under reduced pressure. The crude was triturated in cold MTB to obtain peptide $NH_2$-D-Val-Leu-D-Phe-D-Leu-Leu-OH in sufficient purity to perform the macrocyclization step.

The macrocyclization was adapted from reported procedure[2]. The linear peptide was dissolved in DCM (3 mM), followed by the addition of DIEA (2 equiv.). The reaction mixture was cooled to 0 °C, and HATU (1.5 equiv.) was added. The mixture was then subjected to microwave irradiation at 70 °C for 30 min. After completion, the crude reaction mixture was concentrated under reduced pressure and purified by reverse-phase HPLC (solvent A: water containing 0.05% formic acid (FA); solvent B: acetonitrile (ACN) containing 0.05% FA; gradient 10–80% B over 30 min, held at 80% B for 5 min, then increased to 100% B over 10 min), affording **6** as a white lyophilized powder (27%).

The NMR spectroscopic data were consistent with those previously reported[2,26].

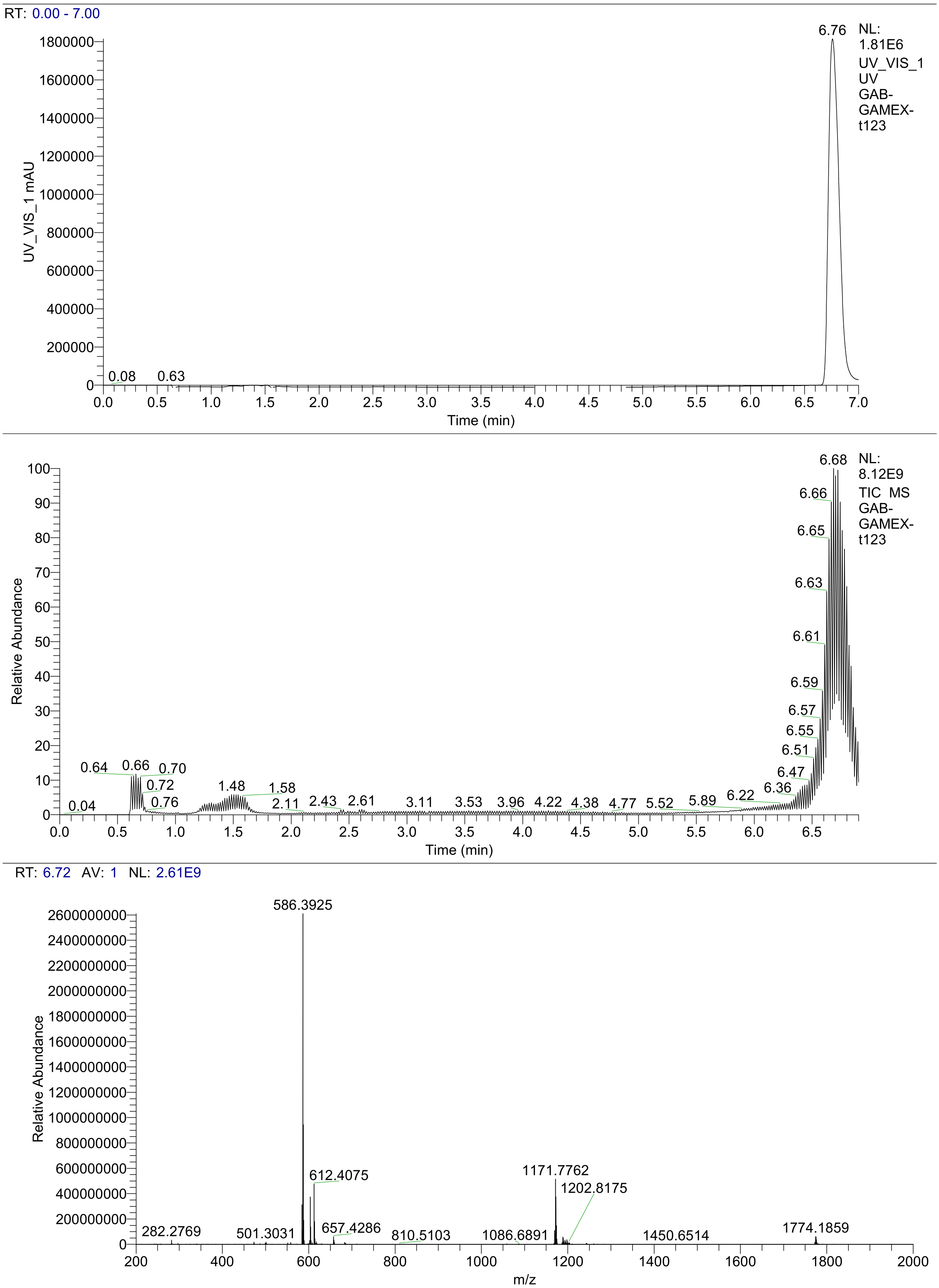

**Supplementary Figure S15.** HRMS (ESI+) spectrum and UV chromatogram of **6.**

**Biochemical Characterization of AI-2 and WT GxpS_T3 variants**

**Cloning:**

The genes encoding the proteins WT-ATC, WT-AT, WT-TC, and WT GxpS_T3 were amplified from the plasmid pJW76_pCOLA_ara_tacI_II_GxpS using the primer pairs 1/2, 1/3, 4/2, and 4/3, respectively. Corresponding AI-modified variants (AI-ATC, AI-AT, AI-TC, and AI-T3) were amplified from the plasmid pJW76_mod_sacB using the primer pairs 1/2, 1/5, 4/2, and 4/5, respectively. The vector backbone was amplified from a pET28a vector with primers 3 and 4. The resulting fragments were assembled by Gibson assembly to generate the final constructs. In each construct, the expressed protein contains an N-terminal hexa-histidine tag followed by a TEV protease site. The exact sequence added at the N-terminal of each protein is, MGSSHHHHHHSSGENLYFQG.

**Expression and solubility test:**

For checking the protein expression, the plasmids were transformed into *E. coli* BL21 Star (DE3) and plated on LB agar containing 30 µg/ml Kanamycin, followed by overnight at 37 °C. Single colonies were used to inoculate 2 ml secondary cultures (1% inoculum), which were grown in glass tubes at 37 °C and 130 rpm, until the $OD_{600}$ reached 0.6-0.8. Cultures were then cooled to 18 °C and induced with isopropyl-β-d-thiogalactopyranoside (IPTG) at final concentrations of 0.02, 0.05, 0.1 and 0.2 mM and grown further for 16 h at 18 °C and 130 rpm. The $OD_{600}$ values of induced and uninduced samples were normalized prior to analysis of protein expression using SDS-PAGE.

Protein solubility was assessed using a similar protocol. Briefly, 50 ml of cultures were induced with 0.05 mM IPTG and grown for 16 h at 18 °C and 130 rpm. Cells were harvested by centrifugation and resuspended in an ice-cold lysis buffer (buffer A) containing 50 mM Tris pH 8.0, 300 mM NaCl, 5% glycerol. Cell densities were normalized, and lysis was performed by sonication (BANDELIN SONOPLUS; MS72 microtip probe; at 50% Amplitude with pulse of 1 sec on and 8 sec off and total 15 min). Unlysed cells were removed by centrifugation at 6,000 × g for 10 min and the remaining lysate was centrifuged at 20,000 × g. The resulting supernatant (soluble fraction) and pellet (insoluble fraction) were separated and analyzed by SDS-PAGE.

**Protein expression and purification:**

Large-scale purification of AI-T3 was performed from the soluble fraction, whereas WT GxpS_T3 was purified from the insoluble fraction. 2 liters of LB broth supplemented with 30 µg/ml Kanamycin was inoculated with a 1% (v/v) overnight *E. coli* culture and grown at 37 °C with shaking at 130 rpm to an $OD_{600}$ of approximately 0.8. Protein expression was induced with IPTG, followed by harvesting at 6000 × g for 10 min and storage of cell pellets at -20 °C until further use.

For AI-T3, expression was induced with 0.2 mM IPTG and cultures were incubated at 18 °C and 130 rpm for 16 h. Thawed cell pellets were resuspended in 40 ml of ice-cold buffer A supplemented with a tablet of protease inhibitors (cOmplete™ EDTA-free Protease-Inhibitor-Cocktail, Roche). Cells were lysed by sonication (BANDELIN SONOPLUS, TS 113 probe) at 58% Amplitude using 1 s on/8 s off pulses for a total time of 30 min. Cell debris were removed by centrifugation at 30,000 g for 1 h. The clarified supernatant was supplemented with 20 mM imidazole and incubated with 2 ml of HisPure™ Ni-NTA resin (Thermo Fischer Scientific) for 30 min with gentle mixing. The resins were washed with 200 ml of buffer A containing 20 mM Imidazole, and bound protein was eluted using buffer A supplemented with 300 mM imidazole. Eluted fractions were concentrated by 10 kDa molecular weight cut-off Amicon® Ultra

Centrifugal Filters (Merck) and further purified using size-exclusion chromatography (SEC) on a Superdex 200 Increase 100/30 GL column with buffer A. Peak fractions from SEC were pooled, concentrated, flash frozen in liquid nitrogen and stored at -80 °C until further use.

For WT GxpS_T3 expression, expression was induced with 1 mM IPTG followed by incubation at 37 °C and 130 rpm for 3 h. Cell lysis was performed as described above. Insoluble material was collected by centrifugation at 30,000 × g for 1 h and the proteins were extracted using 20 ml buffer A supplemented with 8 M urea with gentle shaking for 2 h at room temperature (RT). The extracted proteins were clarified by centrifugation at 30,000 × g for 1 h. Subsequent purification steps were identical to those used for AI-T3 except all the buffers additionally contained 6 M urea. Eluted fractions were concentrated and stored at RT until further use.

**Protein refolding:**

The purified proteins were refolded using either a rapid or a gradual refolding approach. Rapid refolding was achieved by immediate removal of urea through SEC, in which protein was loaded onto a Superdex 200 Increase 100/30 GL column equilibrated with buffer A. In contrast, slow refolding was performed by stepwise dialysis. For this method, the eluted protein, initially in a buffer composed of 50 mM Tris pH 8.0, 300 mM NaCl, 5% glycerol, 300 mM imidazole and 6M urea, was dialyzed sequentially against buffer containing 50 mM Tris pH 8.0, 300 mM NaCl, 5% glycerol, 0.4 M L-Arginine, and decreasing concentrations of urea (4 M, 2 M, 1 M, and 0 M). Following dialysis, the protein was concentrated and subjected to SEC with buffer A.

**Thermal denaturation and melting temperature (Tm) determination:**

Protein samples were prepared at a final concentration of approximately 30-40 µM in buffer A. Samples were centrifuged at 20,000 × g for 15 min to remove particulates prior to measurements. Aliquots of approximately 10 µL were loaded into the capillaries (Prometheus standard capillaries, Nanotemper), ensuring the absence of air bubbles.

Thermal denaturation experiments were performed using a Prometheus Panta nanoDSF instrument (NanoTemper Technologies, 2023) controlled by Panta control software (v1.9). Intrinsic protein fluorescence was monitored as a function of temperature, measuring emission at 330 nm and 350 nm, upon excitation at 280 nm. Samples were heated from 25 °C to 100 °C with a constant heating rate of 3.0 °C/min. Fluorescence signals were recorded continuously throughout the temperature ramp. Each condition was measured in three technical replicates.

Thermal unfolding curves were analyzed using Panta analysis software (v1.8). The ratio of fluorescence intensities (350nm/330nm) was calculated and plotted as a function of temperature. The apparent Tm was determined as the inflection point of the unfolding transition, obtained by the first derivative curve. Data are presented as mean ± SD of three biological replicates where each biological replicate represents the mean of three technical measurements.

**List of primers:**

1. 5'-GAGAATCTTTATTTTCAGGGCGTATGTGTCCATCAGTTGTTTG-3'
2. 5'-GTTAGCAGCCGGATCTCACGCCTGCACCAAACTTGC-3'
3. 5'-GTTAGCAGCCGGATCTCACAAAGATTGGGCCAATAC-3'
4. 5'-GAGAATCTTTATTTTCAGGGCGGTTTCCGCATTGAGCCG-3'
5. 5’-GTTAGCAGCCGGATCTCATAACCGTGTCGCCAGTTC-3’
6. 5’-TGAGATCCGGCTGCTAACAAAG-3’
7. 5’-GCCCTGAAAATAAAGATTCTCGCC-3’
